# Multidimensional control of ingestive behavior by lateral hypothalamic neurotensin neurons

**DOI:** 10.64898/2026.07.17.737325

**Authors:** Dustin Sumarli, Mary C. Loveless, Grace O. Davis, Kyle W. Schroeder, Siyi Ma, Caeley L. Bryan, Emily Vu, McKenna Kernan, Adam Gordon, Garret D. Stuber, Gregory J. Morton, Marta E. Soden

**Affiliations:** Department of Pharmacology, University of Washington, Seattle, United States; Graduate Program in Neuroscience, University of Washington, Seattle, United States; University of Washington Medicine Diabetes Institute, Department of Medicine, Seattle, United States; Department of Anesthesiology and Pain Medicine, University of Washington, Seattle, United States

## Abstract

Consumption of food and water is regulated by interactions between neural circuits that govern motivational drive states, arousal, motor activity, and reward signaling, among other factors. Neurons in the lateral hypothalamus that express the neuropeptide neurotensin (LH-Nts neurons) are known to influence many of these elements, but their precise role in regulating specific aspects of ingestive behavior remains unclear. Utilizing a tightly controlled head-fixed task, we find that LH-Nts neurons strongly encode the rate of licking for water and sweet solutions, with weaker modulation of activity by solution identity and restriction state. Silencing LH-Nts neurons reduces water intake but has little direct effect on hunger or satiety. Instead, we find that these neurons impact multiple underlying behavioral components necessary for food consumption, including arousal and engagement with a novel food source, and also influence thermoregulation and metabolism. Together, these data establish tonic activity of LH-Nts neurons as a critical signal supporting exploration and volitional movement, while also delineating a role for these neurons in driving active consumption, particularly of water. Furthermore, quantitative projection mapping revealed widespread innervation of structures linked to thirst, arousal, reward, metabolism, and facial motor control, suggesting that these neurons play an important role coordinating neural circuits governing multiple different aspects of ingestion.

## Introduction

The process of seeking, finding, and consuming food and water requires the integration of internal drives, such as hunger and thirst, with behavioral modulators, such as arousal and reward learning, to generate specific motivated actions^1–4^. Many neural circuits in the brain contribute to one or more of these processes to various degrees. However, neurons in the lateral hypothalamus (LH) that express the peptide neurotensin (Nts), have been linked to nearly all of the modalities required for feeding and drinking, including motor control, arousal, motivation, reward signaling, and energy balance^5–9^. Despite multiple lines of evidence pointing to an important role for this subpopulation, how the specific activation patterns of LH-Nts neurons might drive or coordinate different components required for ingestive behavior remains largely unknown.

We previously found that optogenetic activation of LH-Nts neurons is strongly reinforcing, an effect driven by activation of dopamine neurons in the ventral tegmental area (VTA)^10^. Several other studies have found that chemogenetic activation of LH-Nts neurons promotes water intake, locomotor activity, wakefulness, and hyperthermia^9,11–15^. However, inconsistent effects have been reported on food intake, with studies reporting decreases^13,15^, no change^12,14^, increases^11^, and increases of high fat diet intake only^13^. These results are likely complicated by the fact that LH-Nts stimulation strongly activates dopamine neurons^10,14^, and reductions in food intake may be a result of a hyperdopaminergic state and resultant hyperactivity, as has been seen following injection of amphetamine or dopamine receptor agonists^16^.

Previous data indicate that the activity of LH-Nts neurons is tied to food and water consumption. Elevated Fos protein has been detected in LH-Nts neurons in dehydrated mice^17^, and single cell calcium imaging of these neurons identified excited and inhibited populations when a mouse entered a section of an arena containing either food or water, though signals were not specifically time-locked to consumption^11^. We previously measured calcium activity in LH-Nts neurons using photometry and found large increases during retrieval and consumption of a sucrsoe reward pellet^10^. In addition to consumption-related activation, photometric recordings of LH-Nts neurons have shown increased activity during wakefulness and REM sleep compared to NREM sleep, indicating a potential role in arousal^18^.

While these studies together support a role for LH-Nts neurons in food and water intake, it remains unclear how the activity of these neurons may reflect reward value in the context of varying internal drive states, or how the regulation of multiple underlying behavioral components (i.e. arousal, motivation, thirst) by LH-Nts signaling ultimately influences rates of consumption. Here we used multiple approaches to investigate the activity patterns and functional output of LH-Nts neurons across different behavioral domains. We utilized a precisely controlled head-fixed licking task that allowed us to determine the contributions of internal state, consumption, and reward value to LH-Nts neuron activation. We also established the impact of silencing these neurons on behaviors including consumption, reward learning, and volitional and reflexive movement, informing a better understanding of the broad functional role of LH-Nts signaling.

We find that LH-Nts neurons respond to the act of licking, with the calcium signal largely driven by lick rate but also influenced by restriction state and solution identity, with the strongest correlated activity associated with water rather than sucrose consumption. LH-Nts neurons are also activated during solid food consumption in an operant task, with the signal enhanced by novelty. In freely moving mice, LH-Nts activity is correlated with movement velocity, and chronic silencing of LH-Nts neurons reduces volitional movement with little effect on reflexive movement. Silencing LH-Nts neurons also reduces water intake, body weight, and body temperature, and impairs engagement with novel food sources and operant responding for food reward without changing overall food intake. Performing a quantitative analysis of the axonal projections of LH-Nts neurons, we identified broad innervation of many brain regions linked to arousal, motivation, thirst, and facial motor control. We conclude that LH-Nts neurons simultaneously impact several underlying behavioral domains necessary for motivated reward seeking and consumption, potentially providing a coordinating signal across multiple neural networks to facilitate ingestive behavior.

## Results

### LH-Nts neurons strongly encode lick rate and are modified by solution identity and restriction state

In order to precisely investigate the activation patterns of LH-Nts neurons during consumption, we injected Nts^Cre^ mice in the LH with AAV-FLEX-GCaMP6m and implanted a fiber optic for photometric recording (**Figure 1A-B**). We utilized a head-fixed multispout setup^19^ in which mice are presented with up to 5 solutions from a retracting and rotating spout, and each lick during a 3 second access period triggers the dispensing of a fixed volume of liquid (**Figure 1C**). Following initial training, water restricted mice received 5 daily sessions in which they were randomly presented with spouts dispensing water, 10% sucrose, 20% sucrose, 20 mM saccharin (noncaloric sweetener), or a dry spout (12 trials per spout, 60 total trials per session, 15-20 s ITI). Mice licked more robustly for all solutions than for a dry spout, with increased licking to sweet solutions over water (**Figure 1D-F**, data averaged from sessions 3-5). Mice decreased licking for all solutions over the course of the session as they became satiated (**Figure 1E**).

**Figure 1.**
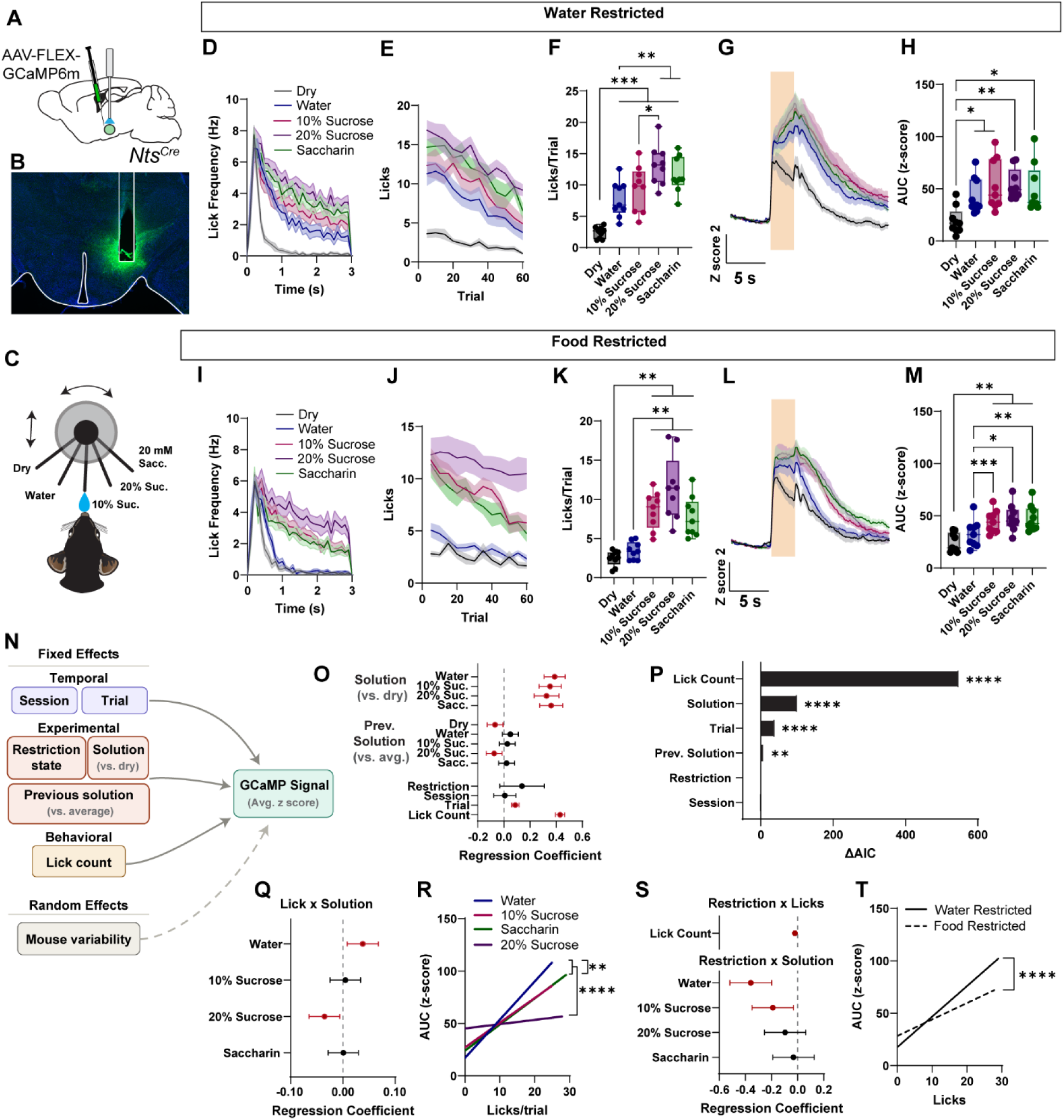
LH-Nts neuron activation during licking is primarily driven by lick count. **A)** Schematic of AAV-FLEX-GCaMP6m injection and fiber optic implantation into the LH of Nts^Cre^ mice. **B)** Example histology image. **C)** Schematic of head-fixed multispout setup. **D)** Lick frequency under water restriction for each solution across the 3 s access period. **E)** Licks per trial for each solution over the course of the session. **F)** Average licks per trial for each solution. (One-way RM ANOVA F=50.13 P<0.0001, Tukey’s multiple comparisons **p<0.01, ***p<0.001.) **G)** Z-scored GCaMP photometry traces during licking. Shaded bar indicates period of spout availability. **H)** AUC of the photometry signal from 0-10 s following spout extension. (One-way RM ANAOVA F=11.25 P=0.0006, Tukey’s multiple comparisons *p<0.05, **p<0.01.) For D-H: N=9 mice, data averaged across days 3-5 of multispout training under water restriction. **I)** Lick frequency under food restriction for each solution across the 3 s access period. **J)** Licks per trial for each solution over the course of the session. **K)** Average licks per trial for each solution. (One-way RM ANOVA F=31.87 P<0.0001, Tukey’s multiple comparisons **p<0.01, ***p<0.001, ****p<0.0001.) **L)** Z-scored photometry traces during licking. Shaded bar indicates period of spout availability. **M)** AUC of the photometry signal from 0-10 s following spout extension. (One-way RM ANOVA F=16.35 P<0.0001, Tukey’s multiple comparisons *p<0.05, **p<0.01.) For I-M: N=9 mice, data averaged across days 3-5 of multispout training under food restriction. **N)** Schematic of mixed effects linear model for photometry data analysis. **O)** Regression coefficients calculated by the linear model. For coefficients highlighted in red, P<0.05, Wald test. **P)** Change in AIC metric for each variable, determined by dropping each term individually and comparing to the full model. Likelihood ratio test, ****P<0.0001, **P<0.01. **Q)** Regression coefficients calculated for a lick x solution interaction term added to the model. For coefficients highlighted in red, P<0.05, Wald test. **R)** Linear regression of individual trial lick count versus AUC of the photometry signal, separated by solution. See Figure S1 for plots with individual data points. Comparison of slopes, **P<0.01, water vs. 10% sucrose or saccharin, ****P<0.0001, 20% sucrose vs. all other solutions. **S)** Top: Regression coefficient calculated for a restriction state x lick count interaction term added to the model. Bottom: Regression coefficients calculated for a restriction state x solution interaction term added to the model. For coefficients highlighted in red, P<0.05, Wald test. **T)** Linear regression of individual trial lick count versus AUC of the photometry signal, separated by restriction state. See Figure S1 for plots with individual data points. Comparison of slopes, ****P<0.0001. Data in D, E, G, I, J and L are presented as mean ± SEM. Box and whisker plots depict the median, 25^th^ and 75^th^ percentiles (box) and min to max (whiskers). Coefficient values are presented with 95% confidence intervals.

Presentation of the dry spout caused an increase in calcium activity in LH-Nts neurons, which was maintained for the spout extension period before returning to baseline (**Figure 1G-H**). A small fluctuation in the calcium signal upon spout retraction was consistently seen throughout the experiment, likely triggered by the sound and motion of spout retraction. For all solutions presented, we observed increases in the calcium signal above the dry spout level during the combined lick and post-lick periods (0-10 s following spout extension), quantified as area under the curve (AUC) of the z score, but there were no significant differences in the GCaMP signal between solutions (**Figure 1G-H**).

Next, mice were switched from water restriction to food restriction and received 5 additional testing days. Under food restriction, mice licked for sweet solutions significantly more than for water or for the dry spout (**Figure 1I-K**). Licking for 20% sucrose was maintained throughout the session, while licking for 10% sucrose or saccharin decreased over the course of the session (**Figure 1J**). Comparing licks under water versus food restriction, we found that mice under food restriction licked significantly less for non-caloric solutions (water and saccharin) than mice under water restriction, but licking for 10% or 20% sucrose was not different between conditions (**Figure S1A**). Under food restriction we again saw a robust increase in calcium signal during the spout extension period for all spouts (**Figure 1L**). The AUC was significantly greater during licking for sweet solutions compared to the water or dry spout (**Figure 1M**).

Because we observed sustained activation of LH-Nts neurons during presentation of the dry spout, we asked whether this activation was dependent on the mouse licking to sample the spout. Across both restriction states we isolated dry spout trials and compared those with zero licks to those with at least one lick. We found that the increased calcium signal was present even in no lick trials, but the AUC was significantly higher in lick trials, indicating that there are both lick-dependent and lick-independent components of the photometry signal (**Figure S1B-C**).

In order to disentangle the impact of different variables on the calcium signal, we generated a linear mixed effects model accounting for restriction state, solution, solution of the immediately preceding trial, number of licks, session number, and trial number (**Figure 1N**), and ran this model on the data from all multispout sessions, using an average of the z-scored calcium signal from 0-10 s following spout extension. Solution was treated as a categorical variable and was compared to the dry spout baseline. In this combined dataset, the regression coefficient for each solution was significant compared to dry spout, consistent with our AUC analysis (**Figure 1O**).

Analyzing the impact of the previous trial solution on the current trial signal, we found small but significant negative regression coefficients when the previous spout was either dry or 20% sucrose, indicating that an immediate history of either no solution or the maximum sucrose concentration had a small negative impact on the calcium signal of the current trial (**Figure 1O**). Regression coefficients for restriction state (food restricted relative to water restricted) and overall session number were not significant, but the model identified a significant positive effect of trial number and lick count on the calcium signal (**Figure 1O**).

To determine which variable had the greatest impact on the calcium signal, we ran repeated iterations of the model, omitting one variable each time. The change in the Akaike Information Criterion metric (ΔAIC) between the reduced and full model served as a measure of each variable’s contribution. Removing lick count, solution identity, trial number, or previous solution each significantly reduced the model fit, as determined by likelihood ratio test, with the largest effect seen with lick count (**Figure 1P**). This indicates that the number of licks made (or, relatedly, the volume of liquid consumed) is the biggest predictor of the calcium signal, rather than the solution identity, restriction state, or trial number within the session.

We also generated alternative versions of the model, adding terms to ask if there was an interaction between the effects of different predictors on the calcium signal. First, looking for an interaction between lick count and solution identity, we found that the effect of lick count on the calcium signal was positive for water trials, and negative for 20% sucrose trials, relative to the dry spout control (**Figure 1Q**). Consistent with this finding, plotting a linear regression of lick count versus AUC of the calcium signal separated by solution, we found a significantly steeper correlation for water trials than for 20% sucrose trials (**Figure 1R and Figure S1D-G**).

Next we asked if there was an interaction between restriction state and lick count, or restriction state and solution identity. We found a small but significant negative effect of food restriction relative to water restriction on the impact of lick count on the calcium signal (**Figure 1S**). Consistent with this, a linear regression of lick count versus AUC for all trial types revealed a significantly steeper correlation in water restricted compared with food restricted mice (**Figure 1T and Figure S1H-I**). Separating by solution, we found that food restriction had a significant negative effect on the impact of water and 10% sucrose trials, but not on saccharin or 20% sucrose trials (**Figure 1S**).

Together, these data show that there is substantial activation of LH-Nts neurons when presented with the potential for consumption (spout extension), even when no liquid is available, with additional activation of these neurons largely dependent on the number of licks in a given trial. Beyond the large effect of lick count, we found that the calcium signal per lick is smaller in food restricted compared to water restricted mice, and that water has a more positive impact per lick than sweet solutions.

### Silencing LH-Nts neurons impairs liquid consumption in a state-dependent manner

Our analysis suggests that LH-Nts neurons preferentially encode water consumption over liquid sucrose consumption. To test the functional impact of LH-Nts neurons on licking for water and sucrose in the head-fixed multispout task, we injected Nts^Cre^ mice bilaterally in the LH with a virus encoding the light chain of tetanus toxin (AAV-FLEX-TeTox-GFP), which prevents neurotransmitter and neuropeptide release (**Figure S2A-B**). Water restricted Nts-TeTox and control mice were trained as described above, and performance was averaged over days 3-5 of testing. Nts-TeTox mice exhibited significantly fewer licks for all solutions compared to control, particularly during the early trials of the session (**Figure 2A-B and S2C**).

**Figure 2.**
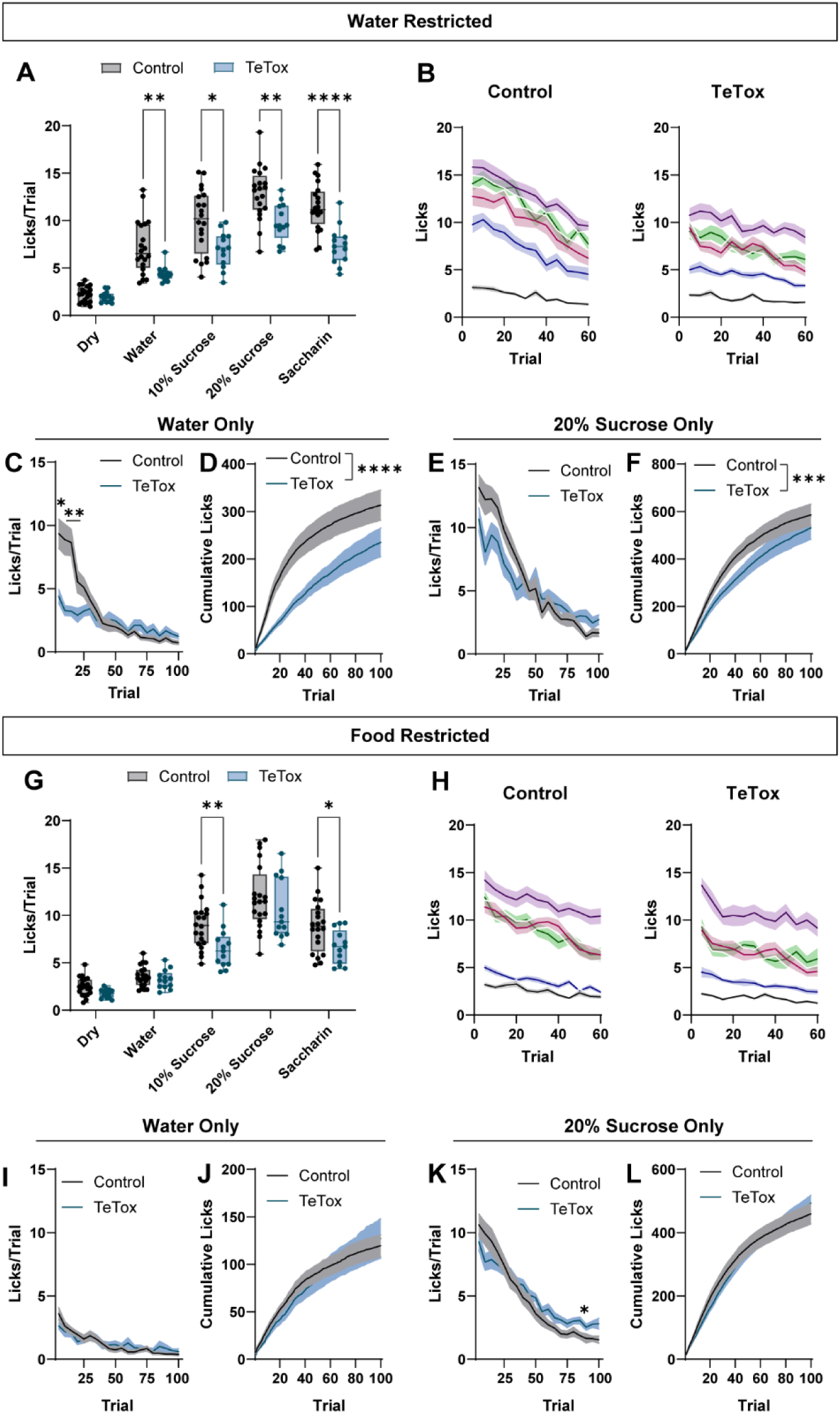
Silencing LH-Nts neurons impairs licking for water and sucrose in a state-dependent manner. **A)** Average licks per trial under water restriction for each solution for control mice and mice expressing TeTox in LH-Nts neurons. (2-way RM ANOVA F_(3.423, 106.1)_=6.342 P=0.0003, Sidak’s multiple comparisons *p<0.05, **p<0.01, ****p<0.0001. Data averaged across days 3-5 of training. N=20 control, 13 TeTox mice.) **B)** Licks per trial for each solution over the course of the session for control and TeTox mice under water restriction. **C)** Licks per trial (averaged in 5 trial bins) for the water only session (2-way ANOVA F_(3.996, 119.9)_=9.688, P<0.0001, Sidak’s multiple comparisons, *p<0.05 bin 1, **p<0.01 bins 2-3). **D)** Cumulative licks over the course of the session (Kolmogorov-Smirnov (KS) test, P<0.0001). **E)** Licks per trial (averaged in 5 trial bins) for the sucrose only session. (2-way RM ANOVA F_(4.285, 128.6)_=4.910, P=0.0008, no significant post hoc comparisons.) **F)** Cumulative licks over the course of the session (KS test, P<0.001). **G)** Average licks per trial under food restriction for each solution. (2-way RM ANOVA Effect of solution F_(4, 124)_=131.6, P<0.0001, Effect of virus F_(1, 31)_=7.011, P=0.0126, Sidak’s multiple comparisons *p<0.05, **p<0.01. Data averaged across days 3-5 of training). **H)** Licks per trial for each solution over the course of the session under food restriction. **I)** Licks per trial (averaged in 5 trial bins) for the water only session. **J)** Cumulative licks over the course of the session (KS test P=0.4676). **K)** Licks per trial (averaged in 5 trial bins) for the sucrose only session (2-way RM ANOVA F_(4.705, 145.9)_=2.420, P=0.0418, Sidak’s multiple comparisons, *p<0.05). **L)** Cumulative licks over the course of the session (KS test P=0.2809). Box and whisker plots depict the median, 25^th^ and 75^th^ percentiles (box) and min to max (whiskers). All other panels presented as mean ± SEM.

To determine the effects of silencing LH-Nts neurons on satiety, water restricted mice were given a single session in which all 5 spouts delivered water for a total of 100 trials. As expected, control mice exhibited high levels of licking on initial trials, but reduced licking as they became satiated, while Nts-TeTox mice licked at low levels throughout the session, as illustrated by the number of licks per trial (**Figure 2C**) and the cumulative licks across the session (**Figure 2D**). To test whether this effect was specific for water, on a separate day, mice were tested with 20% sucrose in all 5 spouts. There was no significant difference in licks between groups at any time point during the session, but we observed a trend towards decreased licks in the Nts-TeTox group in the first half of the session and increased licks relative to control in the second half of the session (**Figure 2E**), which was reflected in a significant shift in the cumulative lick plot (**Figure 2F**).

In contrast to water restriction, when mice were switched to food restriction and tested with the multi-solution set they showed little licking for water and there was no difference in water licks between groups (**Figure 2G**). Nts-TeTox mice licked less than control mice when the caloric content of the solution was low (10% sucrose) or zero (saccharin), but there was no significant difference in licks for the solution with the highest caloric content (20% sucrose) (**Figure 2G-H and S2D**). When tested with water in all 5 spouts, food restricted mice showed low licking and there was no difference between groups (**Figure 2I-J**). When tested with 20% sucrose in all spouts, we observed modestly reduced licking early in the session in Nts-TeTox mice with sustained licking above control levels later in the session; however, the cumulative lick histograms were not significantly different (**Figure 2K-L**).

### Silencing LH-Nts neurons profoundly impacts water intake and energy balance

Our results indicate that silencing LH-Nts neurons in hungry mice reduces consumption of no-or low-calorie solutions, but has a minimal effect when the solution has high caloric value. These findings suggest that LH-Nts neurons do play some role in modulating consumption in a manner that is sensitive to energy demands. To investigate this further, we assessed food consumption and metabolism in mice expressing TeTox or YFP control in LH-Nts neurons (**Figure 3A**). Approximately 6 weeks following viral injection we saw a reduction in body weight as a percentage of the pre-surgery weight in TeTox mice (**Figure 3B and S3A**). This corresponded with a decrease in fat mass but no change in lean mass (**Figure 3C-D and S3B**). Both male and female mice were tested, and similar effects were seen in both sexes. For body composition and metabolic measurements, results in the main figure are pooled across sex, and sex-separated data are presented in supplementary data for metrics that differed by sex in control animals.

**Figure 3.**
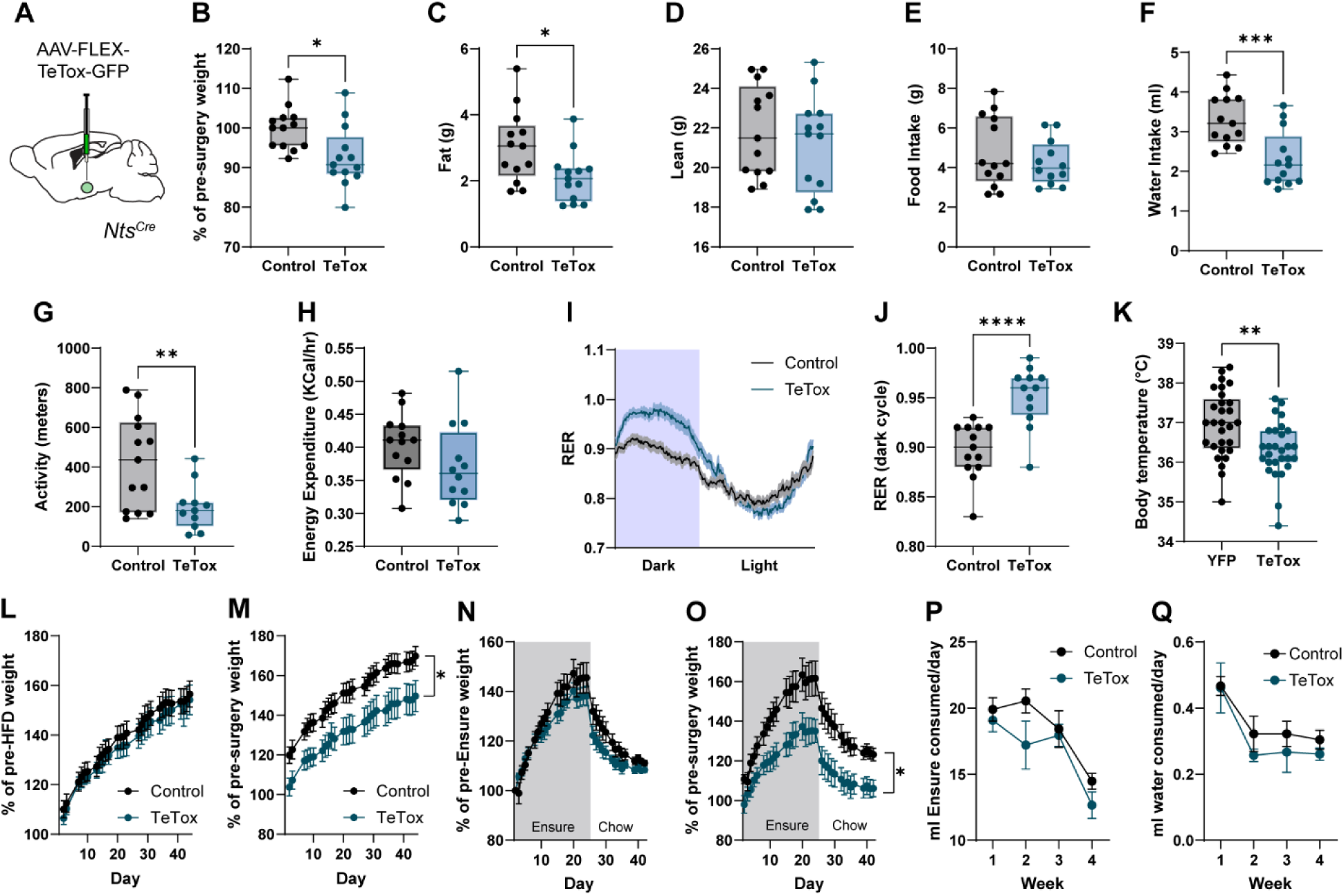
Silencing LH-Nts neurons impairs water intake and alters energy balance. **A)** Schematic of AAV-FLEX-TeTox-GFP injection (bilateral) into the LH of Nts^Cre^ mice. **B)** Percent of pre-surgery body weight 6 weeks following viral injection (N=13 mice/group for panels B-J, Unpaired t test P=0.0107). **C)** Fat mass (Unpaired t test P=0.0197). **D)** Lean mass (Unpaired t test P=0.4627). **E)** 24 hour food intake (Unpaired t test P=0.3790). **F)** 24 hour water intake (Unpaired t test P=0.0107). **G)** 24 hour total activity (Unpaired t test P=0.0078). **H)** Energy expenditure (KCal/hr, averaged over 24 hours; Unpaired t test P=0.1738). **I)** Respiratory exchange ratio across 24 hours in 5 min bins. **J)** Average RER during the dark cycle (Unpaired t test P<0.0001). **K)** Core body temperature (N=29 control, 28 TeTox; unpaired t test P=0.0022). **L)** Body weight on high fat diet as a percent of weight on chow diet (N=12 control, 14 TeTox). **M)** Body weight on high fat diet as a percent of pre-virus injection weight (2-way RM ANOVA, Effect of virus: F_(1, 23)_= 5.436, P=0.0288). **N)** Body weight on Ensure diet, then after return to chow diet, as a percent of pre-Ensure weight on chow diet (N=11 Control, 12 TeTox). **O)** Body weight on Ensure diet as a percent of pre-virus injection weight (2-way RM ANOVA, Effect of virus: F_(1, 21)_= 7.079, P=0.0146). **P)** Ensure consumed per day across 4 weeks of Ensure diet. **Q)** Water consumed per day across 4 weeks of Ensure diet. Box and whisker plots depict the median, 25^th^ and 75^th^ percentiles (box) and min to max (whiskers). All other panels presented as mean ± SEM.

To precisely measure food and water intake, locomotion, and respiration, mice were placed into calorimetry chambers for at least 72 hours. Although Nts-TeTox mice had decreased body weight, we saw no change in total food intake or in meal size, meal duration, or meal number across the light or dark cycle (**Figure 3E and S3C-E**). Nts-TeTox mice drank significantly less water compared to control mice (**Figure 3F**), consistent with our multispout results, and displayed signs of chronic mild dehydration, including reduced skin turgor. Nts-TeTox mice also showed a significant reduction in locomotor activity, particularly during the dark cycle (**Figure 3G and S3F-G**).

Nts-TeTox mice are leaner and hypoactive relative to controls, but we did not observe a significant difference in energy expenditure (kCal/hr) between groups (**Figure 3H**). However, we did observe a significant increase in the respiratory exchange ratio (RER) in Nts-TeTox mice during the dark cycle (**Figure 3I-J and S3H**), indicating a shift towards higher carbohydrate utilization. We also observed a significant decrease in core body temperature in Nts-TeTox mice compared to controls (**Figure 3K and S3I**), consistent with a hyperthermia that has been reported following LH-Nts activation^9^.

Given the altered metabolic state and lean phenotype associated with silencing LH-Nts neurons, we asked whether these mice could gain weight to the same extent as controls if given a calorie-dense, high-fat diet (HFD). We found that Nts-TeTox mice on HFD gained weight at a similar rate to controls, relative to their weight before starting HFD (**Figure 3L**), but did not catch up to control mice relative to their pre-surgery body weight (**Figure 3M**). Because Nts-TeTox mice show decreased water consumption, we next asked whether they would consume a highly palatable liquid diet (Ensure) if it was their only source of calories. Mice were provided both Ensure and water in their home cages, but no solid food. Again, Nts-TeTox mice gained weight at a similar rate as controls compared to their pre-liquid diet body weight (**Figure 3N**), but did not catch up to the control mice relative to their pre-surgery body weight (**Figure 3O**). Nts-TeTox mice consumed equal volumes of Ensure compared to control mice, while water consumed per day was minimal and did not differ between groups (**Figure 3P-Q**). After four weeks on an Ensure diet mice were returned to a solid chow diet, and both control and Nts-TeTox mice rapidly lost weight at a similar pace (**Figure 3N-O**).

These results are consistent with our previous finding that silencing LH-Nts neurons reduces body weight compared to controls^10^. However, this is in contrast to findings in which ablation of LH-Nts neurons with diptheria toxin (DTA) led to increased body fat and a trend towards increased body weight^20^. Nts neurons are largely distinct from orexin/hypocretin expressing neurons in the LH, but this report observed a decrease in orexin positive neurons in the LH following Nts neuron ablation, suggesting potential collateral toxicity or a downregulation of orexin protein. To test whether expression of TeTox in Nts neurons alters orexin levels in the LH, we injected Nts-Cre mice with either AAV-FLEX-TeTox-GFP or a control virus. Following at least 6 weeks of expression, mice were euthanized and sections of the LH were stained for GFP and orexin. We saw no difference in the number of orexin positive neurons between groups (**Figure S3J-K**), indicating that discrepancies between TeTox silencing and DTA ablation may be related to differential effects on neighboring LH-orexin neurons.

### LH-Nts neurons encode the consumption of novel solid food but not satiety state

Though we saw no changes in food intake in Nts-TeTox mice, we had previously observed an increase in the activity of LH-Nts neurons during sucrose pellet retrieval and consumption in an operant reward task^10^. To explore the relationship between LH-Nts activity and solid food consumption in more detail, we trained Nts^Cre^ mice expressing GCaMP6m in the LH on an operant task in which a press on the active lever led to a 3 s delay followed by a 3 s cue (tone+lever light), followed by sucrose pellet delivery (**Figure 4A**). Mice increased active lever presses and the number of pellets earned across 5 days of training (**Figure 4B and S4A**).

**Figure 4.**
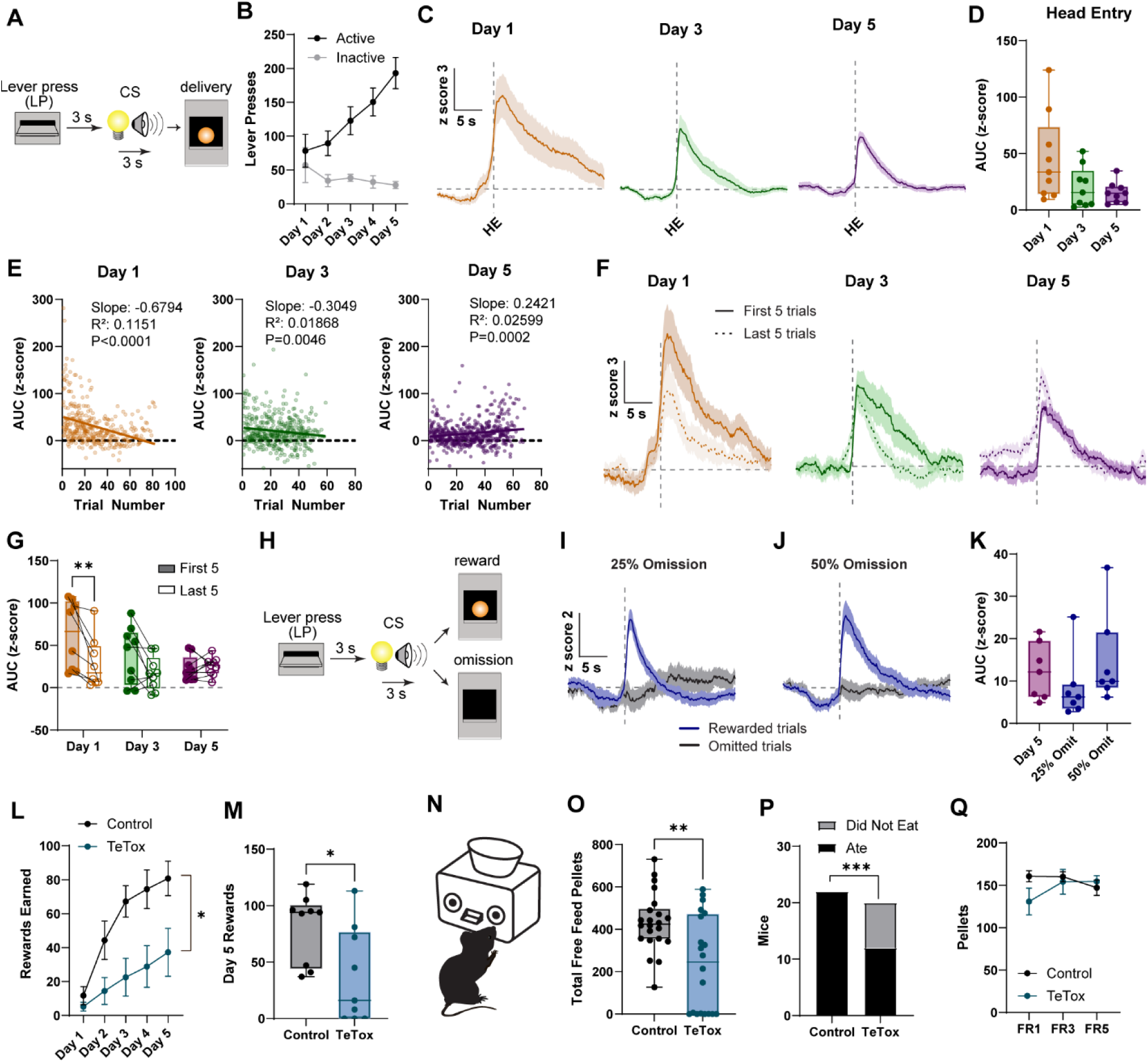
LH-Nts neurons are activated by operant food reward and enhanced by novelty. **A)** Schematic of operant lever press paradigm. **B)** Active and inactive lever presses across training days (N=9 mice for panels B-I). **C)** Average GCaMP signal (peri-event z score) aligned to the first head entry (HE) following pellet delivery across training days. A 4 s baseline period beginning 20 s prior to the head entry was used to calculate the z score. **D)** AUC of the z-scored photometry signal for 0-10 s following head entry (1-way RM ANOVA F_(1.318,10.55)_=4.457, P=0.0516). **E)** Linear regression of trial number versus individual trial AUC across days. **F)** Average GCaMP signal for first and last 5 trials for each mouse across training days. **G)** AUC of the z-scored photometry signal for 0-10 s following head entry for the first and last 5 trials on each day. (2-way RM ANOVA F_(2, 23)_ = 4.140, P=0.0291, Sidak’s multiple comparisons **p<0.01). **H)** Schematic of omission paradigm. **I-J)** Average z-scored photometry traces aligned to first head entry following reward delivery or reward omission during sessions with omission on 25% or 50% of trials. **K)** Average AUC of the z score following head entry during rewarded trials on Day 5 of training or during omission sessions. Due to a technical issue with data collection, 2 mice are excluded from analysis of omission sessions (N=7 mice included, One-way RM ANOVA F_(1.404, 8.423)_=2.423, P=0.1537). **L)** Rewards earned across days for control and TeTox mice in the operant lever press paradigm (N=9 mice/group, 2-way RM ANOVA F_(1.731, 27.69)_=4.309 P=0.0281). **M)** Rewards earned on day 5 (Unpaired t test P=0.0249). **N)** Diagram of FED3 device, with two nose poke ports surrounding a central pellet delivery hopper. **O)** Number of free feed grain pellets consumed over 48 hours of FED3 access in the home cage (N=22 control, 20 TeTox; Unpaired t test P=0.0022). **P)** Number of mice in each group that consumed at least 10 pellets over 48 hours (Fisher’s exact test, P= 0.0011). **Q)** Number of pellets earned in 24 hours on an FR1, FR3, or FR5 nose poke reinforcement schedule. (2-way ANOVA F_(1.791, 57.31)_=2.368 P=0.1082). Box and whisker plots depict the median, 25^th^ and 75^th^ percentiles (box) and min to max (whiskers). All other panels presented as mean ± SEM.

Consistent with our previous results, we observed a small decrease in calcium activity during the delay and cue periods following the lever press on day 1, and a ramping increase in the signal prior to the lever press that developed across training days (**Figure S4B-E**). Aligning the photometry traces to the first head entry into the hopper following pellet delivery (measured by infrared beam break), we found significant increases in calcium activity across all sessions (**Figure 4C**), again consistent with our previous results. Analyzing the AUC for the 10 s following head entry, we observed a trend towards decreasing signal across training days (**Figure 4D**, One-way RM ANOVA p=0.0516).

It is possible that a novelty component amplifies LH-Nts activity on Day 1, or alternatively the calcium response may trend smaller on day 5 because more rewards are consumed, leading to greater satiety. To test these possibilities, we plotted the AUC for each head entry against the trial number for each day. We observed a significant negative correlation between trial number and calcium signal on Day 1 (**Figure 4E**), which was reflected in a decrease in the AUC between the first 5 and last 5 trials of the session (**Figure 4F-G**). However, by Day 5 the correlation was no longer negative (**Figure 4E**), and there was no significant difference in the AUC between the first and last 5 trials on Day 3 or Day 5 (**Figure 4F-G**).

Reductions in LH-Nts neuron activity over learning may also reflect reduced uncertainty, similar to effects seen in dopamine neurons^21^. To test this, trained mice experienced two sessions in which the reward pellet was omitted on either 25% or 50% of trials (**Figure 4H**). The response to the lever press and cue was the same between omitted and non-omitted trials, as the animal could not anticipate whether the reward would be delivered (**Figure S4F**). We saw no GCaMP response when mice made head entries into the hopper following omission trials (**Figure 4I-J**), and the response to retrieval/consumption during rewarded trials was similar in amplitude to what we observed on the last day of FR1 conditioning (100% rewarded; **Figure 4K**). Together, these data support a novelty-related component of the signal during initial training that decreases over time as mice learn the contingency, and not a reward prediction error-like response.

### Silencing LH-Nts neurons impairs operant responding and engagement with novel food

Given the strong activation of LH-Nts neurons during pellet consumption in this task, we next asked whether silencing these neurons impacts operant performance. Control and Nts-TeTox mice experienced 5 sessions of the same lever press task described above. Nts-TeTox mice earned significantly fewer pellets than control mice across training (**Figure 4L**). Examining the distribution of performance during the final session, we found that 3 of 9 Nts-TeTox mice completely failed to engage with the task, making zero lever presses (**Figure 4M**). The impaired acquisition of operant behavior in Nts-TeTox mice could be due to anhedonia, reducing the rewarding value of the sucrose pellet. However, we found no difference between groups in a two-bottle sucrose preference test, though Nts-TeTox mice did consume less total liquid than controls (**Figure S4G-H**).

Another possibility is that the reduced operant performance in Nts-TeTox mice is due to the reduced locomotor activity exhibited by Nts-TeTox mice, which may impair learning during the limited 1 hour sessions. To test whether these mice could acquire operant responding with longer access, mice were singly housed in cages containing FED3 devices^22^, which can deliver pellets in the home cage across a 24 hour period, either noncontingently or in response to a nose poke (**Figure 4N**).

To train mice to use the FED3 we removed all normal chow from the cage and loaded the devices with a grain-based pellet, which would serve as their only source of food. For the first two days, pellets were delivered noncontingently, with a new pellet delivered to the hopper as soon as the previous one was removed. Control mice quickly learned to feed from the FED3 devices, consuming on average over 400 pellets in the course of 48 hours (**Figure 4O**). Notably, 8 of 20 TeTox mice failed to engage with the FED3 device, eating fewer than 10 pellets total over 48 hours (**Figure 4O-P**; mice were monitored and supplemented with a small amount of chow if their body weight approached 85% of ad libitum).

Following two days of free pellet availability, mice were switched to an FR1 contingency such that a single nose poke in the active port was required to trigger pellet delivery. Mice had 24 hours of access to the device on an FR1 schedule, followed by 24 hours at FR3 and 24 hours at FR5. Only Nts-TeTox mice who had learned to eat from the device were put through the FR paradigm, and within this group there was no difference from control in the total number of pellets earned on any day (**Figure 4Q**). Together, these data indicate that, while Nts-TeTox mice will eat normal amounts of chow, a subset are impaired in the acquisition of operant responding for food and in their ability to engage with a novel food source even when an operant response is not required.

### LH-Nts neurons regulate volitional movement with limited impact on reflexive movement

To further examine the impact of LH-Nts neurons on exploration, arousal, and food consumption in a novel environment, mice were fasted for 24 hours and then placed into a novel arena with a single chow pellet secured in the center. Nts-TeTox mice took significantly longer to initiate consumption of the pellet (**Figure 5A**), but there was no difference in total consumption over the 20 minute trial (**Figure 5B**). Nts-TeTox mice also showed significantly reduced locomotor activity in the arena (**Figure 5C-D**), consistent with our results measuring day/night locomotion in the calorimetry chambers. Examining locomotor activity in 5 minute bins over the course of the trial revealed that Nts-TeTox mice lack the conventional exploratory hyperlocomotion displayed by control mice when initially placed into a novel arena (**Figure 5E**).

**Figure 5.**
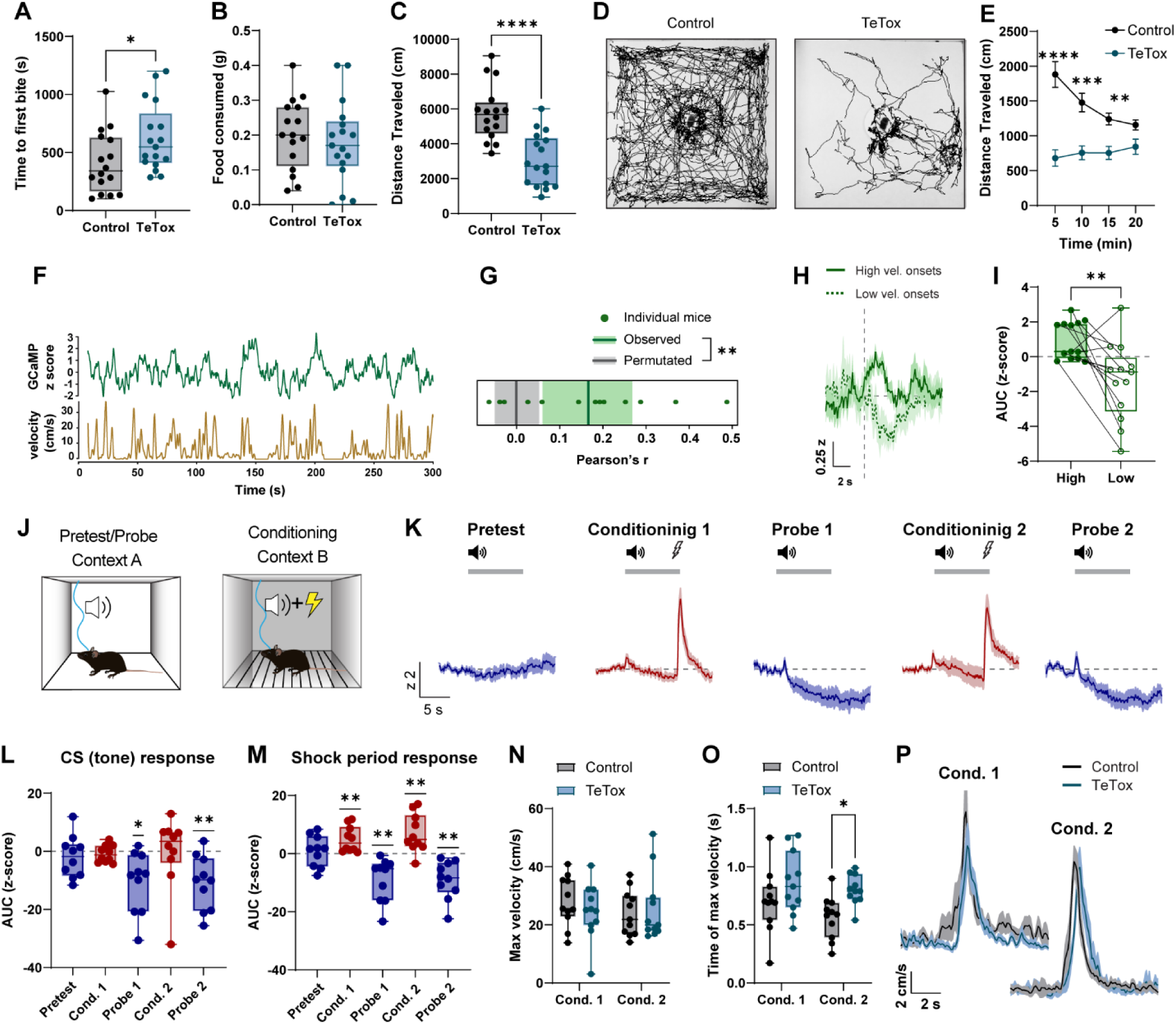
LH-Nts neuron activity correlates with velocity, and silencing LH-Nts neurons impairs volitional but not reflexive movement. **A)** Time to first bite of chow pellet in fast/refeed assay in a novel arena (N=16 control, 17 TeTox, Unpaired t test P=0.0359). **B)** Total grams of chow consumed during the 20 min trial (Unpaired t test P=0.5604). **C)** Distance traveled during the 20 min trial (Unpaired t test P<0.0001). **D)** Example tracks of Control and TeTox mice in the arena. **E)** Distance traveled in 5 minute bins (2-way RM ANOVA, F_(2.217, 68.72)_= 2.82 P<0.0001). **F)** Example trace of GCaMP photometry signal (z-scored to the entire session) and velocity in an open field. **G)** Correlation coefficients of photometry signal and velocity (Pearson’s r) for individual mice (points, N=13) and the mean and 95% CI (green line and shaded region). Gray line and shaded region depicts the mean and CI of 1000 random circular permutations of the velocity trace relative to the photometry trace (permutation test, p=0.002). **H)** Z-scored photometry traces (averaged per mouse) aligned to high and low velocity onset periods (periods when a mouse crossed above the 80^th^ or below the 20^th^ percentile of their velocity range and remained there for at least 2 seconds). **I)** AUC of the average high and low velocity trace per mouse for 5 s following the low or high velocity onset (N=13 mice, paired t test P=0.0073). **J)** Schematic of contexts for Pavlovian fear conditioning. **K)** Z-scored LH-Nts GCaMP photometry traces during presentation of the CS only (Pretest and Probe sessions) or the CS paired with a 0.3 mA footshock (Conditioning sessions). **L)** AUC of the z-scored photometry trace during the 9 s CS presentation (N=10 mice, One-sample t test, *P<0.05, **P<0.01 compared to 0). **M)** AUC of the z-scored photometry trace during the 5 s following shock presentation (One-sample t test, *P<0.05, **P<0.01 compared to 0). **N)** Maximum velocity in response to footshock in Control and TeTox mice. **O)** Time of maximum velocity following shock (2-way RM ANOVA, Effect of session: F_(1, 20)_=5.360, P=0.0313, Effect of virus: F_(1, 20)_=5.742, P=0.0265. Sidak’s multiple comparisons *P<0.05). **P)** Velocity traces in response to shock. Box and whisker plots depict the median, 25^th^ and 75^th^ percentiles (box) and min to max (whiskers). Other panels are presented as mean ± SEM.

To further probe differences in volitional movement, Nts-TeTox and control mice were given access to a running wheel, which is highly motivating in mice^23^. We found a significant reduction in running in Nts-TeTox mice compared to control (**Figure S5A**). Consistent with generalized deficits in volitional movement, we also observed that Nts-TeTox mice show significantly reduced shredding of nesting material (nestlets) within their home cage, quantified using a previously established scale^24^ (**Figure S5B-C**).

Given the robust impairment in locomotor activity that we observed when silencing LH-Nts neurons, as well as previous reports of increased locomotion following activation of these neurons, we asked whether there was a significant correlation between activity of LH-Nts neurons and movement velocity. Nts^Cre^ mice expressing GCaMP6m in the LH were placed into a large open field arena and were allowed to explore for 5 minutes while photometry and video recordings were made. The instantaneous velocity of each mouse was determined via automated tracking software, and photometry and velocity traces were aligned and smoothed (**Figure 5F**). A linear regression was calculated for each animal to measure the correlation coefficient. We then performed a permutation analysis, repeatedly shifting each mouse’s photometry trace relative to the velocity trace by a random amount and calculating the regression for the shifted data. The mean Pearson’s r value (r = 0.166) of the observed data was significantly higher than that of the permutated data (r = -0.001), indicating a meaningful positive correlation between velocity and calcium activity in LH-Nts neurons (**Figure 5G**). For each animal we also calculated the 20^th^ and 80^th^ percentile of all velocity values and identified time points when the animal crossed below the 20^th^ or above the 80^th^ percentile (low and high velocity onsets). Aligning GCaMP signals at these time points, we observed increases in calcium activity following high velocity onsets and decreases following low velocity onsets (**Figure 5H-I**), consistent with our correlation analysis.

Our results are consistent with impairments in volitional movement after LH-Nts silencing; however, we cannot exclude the possibility that these effects are the result of a general motor impairment. To explore this further, we examined reflexive movement in response to a painful stimulus. Mice expressing GCaMP in LH-Nts neurons experienced a Pavlovian fear conditioning paradigm that elicits defensive flight in response to footshock and defensive freezing in response to a conditioned stimulus (CS). Mice were first placed into Context A and exposed to 5 presentations of a 9 s CS tone (Pretest). Mice were then placed into Context B and experienced 10 pairings of the CS with a 0.3 mA co-terminating footshock (Conditioning Session 1). On the second day, mice received a probe session (CS only) in Context A and a second conditioning session in Context B. A final probe session was conducted on the third day (**Figure 5J**).

We saw no activation of LH-Nts neurons to the tone during the pretest (**Figure 5K**). During both conditioning sessions there was minimal activity of LH-Nts neurons during CS presentation, but we observed a strong activation of these neurons to the footshock (**Figure 5K-M**). Notably, during both probe sessions we observed a significant decrease in calcium activity during and following the CS presentation (**Figure 5K-M**). Given the correlation we observed between locomotion and LH-Nts calcium activity, this decrease may be related to cue-induced freezing responses.

To test whether silencing LH-Nts neurons impairs response to footshock, control and Nts-TeTox mice experienced the same Pavlovian fear conditioning paradigm. In both groups we saw an increase in freezing during presentation of the CS during the probe sessions, with no significant difference in freezing between groups (**Figure S5D**). We did not observe a difference in maximum velocity in response to the footshock during conditioning sessions (**Figure 5N**), indicating that LH-TeTox mice retain the ability to respond to a painful stimulus, though there was a slight delay in the time to maximum velocity on the second conditioning day (**Figure 5O-P**).

### LH-Nts neurons innervate brain regions important for thirst, arousal, and motor control

Our fiber photometry and behavioral data implicate LH-Nts neuron function in a multitude of behaviors related to consumption, including thirst, the motor component of licking and eating solid food, volitional movement, general arousal, exploration, engagement with novel food sources, and operant reward learning, as well as thermoregulation and metabolism. The breadth of functions regulated by LH-Nts neurons suggests that they impact many downstream regions throughout the brain. Previous reports have identified some axonal targets of these neurons^9,15^, but a comprehensive projection map had not been generated. In order to better understand how LH-Nts neurons exert influence over such a wide array of behaviors, we conducted an unbiased quantitative analysis of the axonal projections of these neurons throughout the brain.

Nts^Cre^ mice were injected in the LH with a virus encoding synaptophysin tagged with GFP, which directs the fluorescent marker to the axon terminal (**Figure 6A-B**). Serial sections of the brain were imaged and aligned to the Allen Institute Mouse Brain Atlas (**Figure 6C**). The integrated puncta intensity per square micron was calculated for each brain region and normalized within each animal. Regions with an intensity at least 5% of the intensity at the injection site are plotted in **Figure 6D**.

**Figure 6.**
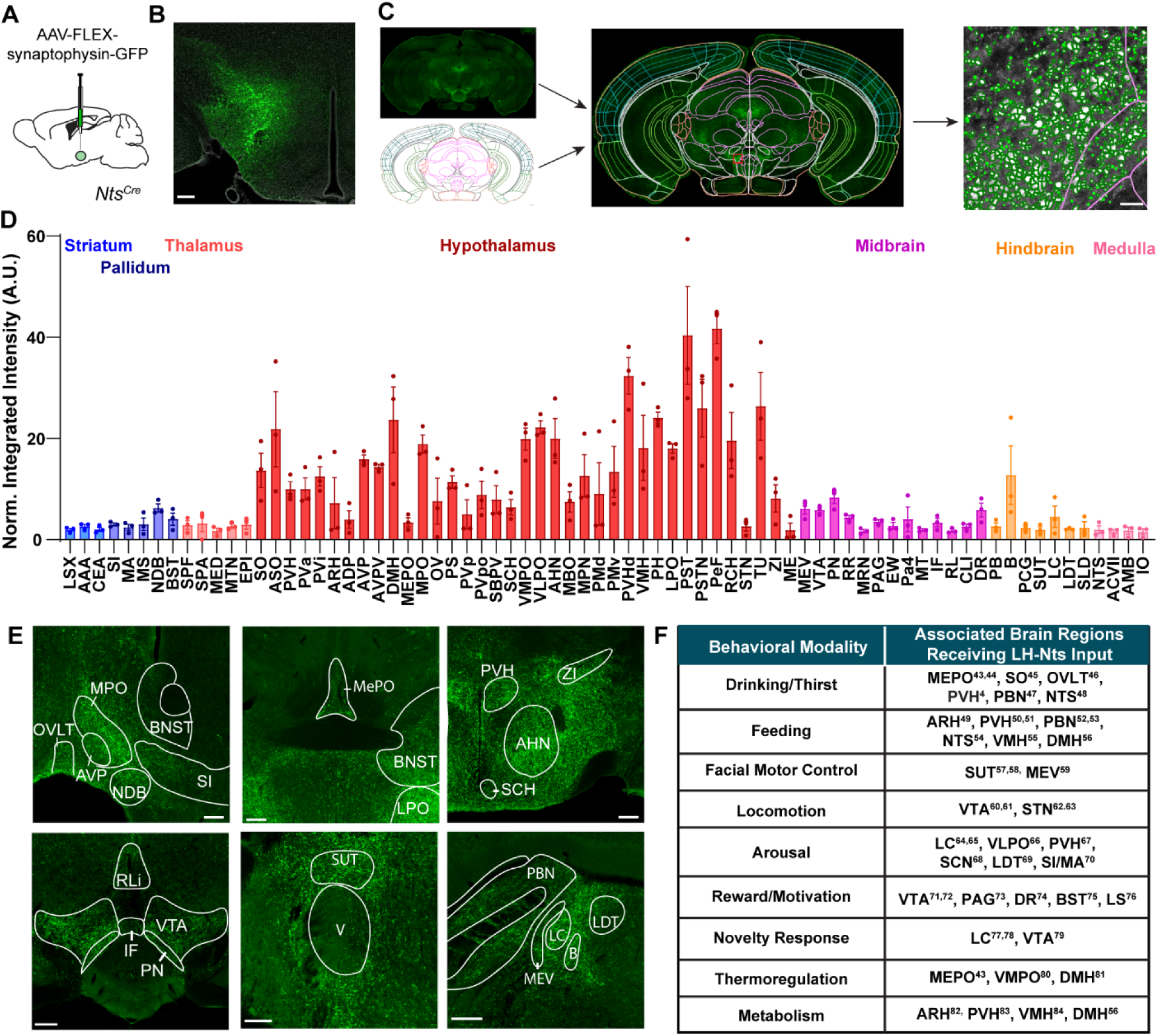
LH-Nts neurons project to target regions associated with impacted behavioral domains. **A)** Schematic of AAV-FLEX-synaptophysin-GFP injection into the LH of Nts^Cre^ mice. **B)** Example image of virus expression in the LH. **C)** Schematic showing alignment of sections with the Allen Brain Atlas and synaptophysin-GFP puncta detection in QuPath. **D)** Normalized integrated puncta intensity across brain regions. Only regions expressing at least 5% of the integrated intensity of the LH are shown. **E)** Example images showing innervation of select regions. **F)** Behavioral modalities linked to LH-Nts neuron activity, and select brain regions known to impact those modalities. Abbreviations: LSX: Lateral septal complex; AAA: Anterior amygdalar area; CEA: Central amygdalar nucleus; SI: Substantia innominate; MA: Magnocellular nucleus; MS: Medial septal nucleus; NDB: Diagonal band nucleus; BST: Bed nuclei of the stria terminalis; SPF: Subparafascicular nucleus; SPA: Subparafascicular area; MED: Medial group of the dorsal thalamus; MTN: Midline group of the dorsal thalamus; EPI: Epithalamus; SO: Supraoptic nucleus; ASO: Accessory supraoptic group; PVH: Paraventricular hypothalamic nucleus; PVa: Paraventricular hypothalamic nucleus, anterior; PVi: Paraventricular hypothalamic nucleus, intermediate; ARH: Arcuate hypothalamic nucleus; ADP: Anterodorsal preoptic nucleus; AVP: Anteroventral preoptic nucleus; AVPV: Anteroventral periventricular nucleus; DMH: Dorsomedial nucleus of the hypothalamus; MEPO: Median preoptic nucleus; MPO: Medial preoptic nucleus; OV: Vascular organ of the lamina terminalis; PS: Parastrial nucleus; PVp: Periventricular hypothalamic nucleus, posterior; PVpo: Pariventricular hypothalamic nucleus, preoptic; SBPV: Subparaventricular zone; SCH: Suprachiasmatic nucleus; VMPO: Ventromedial preoptic nucleus; VLPO: Ventrolateral preoptic nucleus; AHN: Anterior hypothalamic nucleus; MBO: Mammillary body; MPN: Medial preoptic nucleus; PMd: Dorsal premammillary nucleus; PMv: Ventral premammillary nucleus; PVHd: Paraventricular hypothalamic nucleus, descending; VMH: Ventromedial hypothalamic nucleus; PH: Posterior hypothalamic nucleus; LPO: Lateral preoptic area; PST: Preparasubthalamic nucleus; PSTN: Parasubthalamic nucleus; PeF: Perifornical nucleus; RCH: Retrochiasmatic area; STN: Subthalamic nucleus; TU: Tuberal nucleus; ZI: Zona incerta; ME: Median eminence; MEV: Midbrain trigeminal nucleus; VTA: Ventral tegmental area; PN: Paranigral nucleus; RR: Midbrain reticular nucleus, retrorubral area; MRN: Midbrain reticular nucleus; PAG: Periaqueductal gray; EW: Edinger-Westphal nucleus; Pa4: Paratrochlear nucleus; MT: Medial terminal nucleus of the accessory optic tract; IF: Interfascicular nucleus raphe; RL: Rostral linear nucleus raphe; CLI: Central linear nucleus raphe; DR: Dorsal nucleus raphe; PB: Parabrachial nucleus; B: Barrington’s nucleus; PCG: Pontine central gray; SUT: Supratrigeminal nucleus; LC: Locus coeruleus; LDT: Laterodorsal tegmental nucleus; SLD: Sublaterodorsal nucleus; NTS: Nucleus of the solitary tract; ACVII: Accessory facial motor nucleus; AMB: Nucleus ambiguous; IO: Inferior olivary complex.

Consistent with the variety of behaviors that we identified as being regulated by LH-Nts neurons, we saw widespread innervation of subcortical regions throughout the brain that have been linked to relevant behavioral modalities. These areas included structures linked to thirst and fluid intake, such as the vascular organ of the lamina terminals (OVLT) and median preoptic nucleus (MEPO) (**Figure 6D-E**), and hindbrain structures linked to water and food intake, including the nucleus of the solitary tract (NTS) and parabrachial nucleus (PBN). Fibers also innervated structures directly connected to the control of jaw and facial movements, which likely play an important role in the physical act of consumption, including the midbrain trigeminal nucleus (MEV) and the supratrigeminal nucleus (SUT), with fibers encircling the trigeminal motor nucleus (V; **Figure 6E**). We also observed projections to the locus coeruleus (LC) and laterodorsal tegmentum (LDTg), critical arousal centers in the pons. A summary of innervated brain regions that have been linked to behaviors impacted by LH-Nts neurons are noted in **Figure 6F**.

## Discussion

Though LH-Nts neurons have long been linked to multiple behavioral modalities, here we are able to identify for the first time how these neurons encode specific components of consumption, and highlight how they regulate different behavioral elements and metabolic functions to impact food and water intake. LH-Nts neurons are activated by both liquid and solid food consumption, with activation patterns in the head-fixed task most strongly driven by the number of licks, but modulated by the solution identity and the restriction state of the animal, boosting the signal per lick in response to water over sucrose and in thirsty mice compared to hungry mice. Consistent with these results, silencing these neurons impairs water intake but does not impact hunger or satiety for food or sweet solutions. Silencing does, however, impair engagement with a novel food source, consistent with multiple other metrics pointing towards the necessity of these neurons for maintaining general arousal and driving novelty-induced exploration and volitional movement. We also identified an important role for LH-Nts neurons in maintaining energy balance, including by regulating body temperature and controlling the balance of carbohydrate and fat utilization.

Though LH-Nts neurons are a major input to VTA dopamine neurons, activation patterns of these two cell types are different in many ways. In a similar head-fixed task, recordings of dopamine neuron cell bodies and dopamine release demonstrate strong value coding, with scaled responding to increasing sucrose concentrations and negative responses to rewards of low or no value (i.e. when water is offered to food restricted mice)^19,25^. LH-Nts neurons, by contrast show strong activation to spout extension regardless of solution identity or even whether a solution is present at all, and responding that could almost be considered inverse value coding, with the calcium signal per lick higher for water than for sucrose. This pattern is consistent with the clear role for these neurons in driving water intake, but underscores the fact that LH-Nts neurons are just one input of many contributing to patterns of dopamine release.

Some of our results are in contrast to a previous study that used DTA to ablate LH-Nts neurons and found increased as opposed to decreased body weight and fat mass^20^. A likely explanation is the reported impact of ablation on neighboring non-Nts cells, including orexin neurons. Differences in the extent of viral spread or rostral-caudal targeting could also be a factor. Our results are largely consistent, however, with previous reports of LH-Nts neuron chemogenetic activation driving water intake, locomotion, wakefulness, and hyperthermia^9,12–15^. They are also consistent with studies showing reduced locomotion, novelty response, and arousal with chemogenetic inhibition of LH-Nts neurons^9,15,20^.

Previous chemogenetic inhibition studies did not observe a decrease in water intake however, even with chronic CNO administration^15,20^. Silencing with TeTox leads to a more complete and sustained loss of synaptic transmission relative to inhibition with DREADDs, which reduces but does not fully block neuronal activity. This likely accounts for the more robust findings we observed, particularly in domains such as long-term consumption patterns, arousal, and energy balance that are regulated on a time scale of hours and days. Our results highlight the importance of tonic activity of LH-Nts neurons in supporting a number of critical functions.

We found that LH-Nts neurons are activated by a painful footshock and are inhibited during freezing to a shock-predictive cue. However, silencing LH-Nts neurons did not impact acquisition of conditioned freezing and had a minimal impact on locomotor responses to shock. This implies that sensory systems and instinctual defensive responses are largely intact in these animals, and strengthens the argument that these neurons engage a constellation of behavioral elements specifically in support of volitional movement and ingestive behavior.

The role of LH-Nts neurons in promoting thirst and water intake is clear, both in our results and in previous studies. The role for these neurons in regulating food intake appears to be much more nuanced, with prior reports showing both decreases and increases in feeding following LH-Nts activation^11–15^. Our results help to separate a number of different components of food consumption, and support a role for LH-Nts neurons in some aspects of food intake but not others. These neurons do not seem to have a strong role in regulating hunger or satiety. This was reflected in the licking experiments with sucrose only, in which LH-TeTox mice showed somewhat less vigorous licking at the beginning of the session, but maintained licking for longer such that the total volume of sucrose consumed was not different. Consistent with this, in well-trained mice the amplitude of LH-Nts neuron responses showed little change across the course of operant sessions, indicating minimal encoding of satiety state.

Though they play a limited role in control of hunger, LH-Nts neurons impact other underlying behaviors that influence food consumption, including supporting novelty-induced exploration. We not only observed impaired locomotor activity in a novel environment and reduced engagement with a novel food source, but also found that calcium responses in these neurons were larger in the initial trials on the first day of an operant reward task, indicating an enhanced novelty response. Projections of LH-Nts neurons to regions including the VTA and LC may support salience encoding in addition to general arousal, facilitating responding to novel stimuli. Silencing LH-Nts neurons also impaired acquisition of an operant task, leading to reduced reward consumption. This indicates that these neurons may facilitate cue-reward or action-outcome associations, consistent with our previous finding that stimulating LH-Nts to VTA projections is reinforcing^10^. Finally, projections of LH-Nts neurons to regions responsible for facial motor control indicate that these neurons may contribute to regulating the motor patterns that drive chewing and licking. This is consistent with the correlation we observed between lick rate and activation of these neurons, and also consistent with previous observations that stimulating all LH-GABA neurons, of which Nts neurons are a subset, can generate gnawing, chewing, or licking motions even in the absence of food or water^19,26,27^.

We observed complex changes to energy balance and metabolism following silencing of LH-Nts neurons. Though mice had both reduced activity and reduced core body temperature, overall energy expenditure was not significantly altered. However, the combination of hypothermia, reduced fat mass, and elevated dark cycle RER points to a potential dysregulation of both brown and white adipose tissue. Notably, the LH itself, along with other hypothalamic and brainstem regions targeted by LH-Nts neurons, are known to regulate fat utilization and storage^28–30^. In particular, it is known that LH orexin neurons can mediate lipolysis and brown fat neurogenesis^31,32^, and it has been shown that LH-Nts neurons modulate the activity of neighboring orexin cells^20,33,34^. It should be noted that our data are in contrast with the DTA ablation model, in which mice show increased fat mass and reduced overall energy expenditure with no change to RER^20^.

The broad impact of these neurons across multiple distinct behavioral domains is reflected in the axonal innervation of critical downstream regions linked to diverse functions. What remains unknown is the extent to which individual LH-Nts neurons may project to multiple targets, or whether subpopulations of these neurons play specialized roles in regulating different behavioral outcomes. Previous work has found both ascending (to the lateral preoptic area) and descending (to the VTA) projections originating from the same LH-Nts neurons, indicating the potential for widespread collateralization^15^. However, it has also been shown that the subset of LH-Nts neurons expressing the leptin receptor are distinct from the subset that are activated by dehydration, and that the former project to the VTA but the latter do not, indicating genetically distinct and projection-specific subpopulations^17^. Transcriptomic analysis has identified at least two major subsets of LH-Nts neurons, those expressing *Tac1* and those expressing *Crh*^35^, and achieving a more granular division of these neurons is likely with additional analysis of gene expression. Whether these genetically defined subsets of neurons will have non-overlapping projection patterns and functions, or whether individual LH-Nts neurons can simultaneously modulate or coordinate multiple downstream regions via collateralized projections remains to be seen.

## Methods

### Mice

All procedures were approved and conducted in accordance with the guidelines of the Institutional Animal Care and Use Committee of the University of Washington. Mice were group-housed with ad libitum food and water unless otherwise noted. For the majority of experiments mouse were housed on a 12 hour light/dark cycle, with the exception of calorimetry experiments, which were conducted on a 14/10 cycle. Approximately equal numbers of male and female mice between 8 and 20 weeks of age were used for all experiments. *Nts^Cre^* mice (Strain 017525) are available from Jackson Labs.

For all experiments, viral expression and fiber placement were confirmed post-hoc, and mice with improper expression or placement were excluded from analysis.

### Viruses

All viruses were produced in-house by the University of Washington Molecular Genetics Resource Core, as described^36^, or purchased from Addgene. All viruses were serotype AAV1.

### Surgery

Mice were injected at 8-12 weeks of age and recovered for at least 3 weeks prior to experimentation. Mice were anesthetized with isoflurane before and during viral injection and fiber implantation. LH coordinates for injection were (in mm) M-L: ±1.0, A-P: -1.25, D-V: -5.0. For all injections the needle was lowered to 0.5 mm below the indicated depth and slowly raised while the virus was injected. Fiber optics for photometry were 400 μm in diameter (Doric or MBF Bioscience) and were implanted at a depth of 4.9 mm.

### Immunohistochemistry

Mice were perfused with 4% PFA and 50 μm sections were made using a cryostat (Leica) or microtome (RMC). Free-floating sections were stained with primary antibody overnight at 4°C followed by secondary antibody for two hours at room temperature. Primary antibodies used were: Chicken anti-GFP (AbCam 13970, 1:6000), Rabbit anti-Orexin (Sigma-Aldrich AB3704, 1:1000). Secondary antibodies raised in donkey (Jackson ImmunoResearch) were used at a concentration of 1:400. Sections were mounted with DAPI-Fluoromount-G (Southern Biotech) and imaged using a VS200 SlideScanner microscope (Evident Scientific).

### Fiber Photometry

Freely moving recordings were made using an RZ5 Processor and Synapse software (Tucker Davis), with LEDs, filter cubes, and cables from Doric or Thor Labs. Head-fixed recordings were made using an RZ10 Processor (Tucker Davis) with integrated LEDs and filter cubes from Doric. A 465 LED (30-40 μW at the fiber tip) was used to excite GCaMP6m and a 405 LED was used to monitor the isosbestic signal for freely moving recordings only. Fluorescence was returned through the same patch cord, bandpass filtered, and recorded at 1017.25 Hz. See quantification section below for description of photometry signal processing.

For behavioral photometry experiments, signals were aligned to behavioral events either via TTL delivery from MedAssociates software (operant responding and fear conditioning), TTL delivery from the Arduino controlling the head-fixed setup, or by video recording behavior and identifying event times using Ethovision software (Noldus).

### Behavioral Assays

*Head-fixed multispout licking:* OHRBETS head-fixed behavioral systems were manufactured in house as described^19^. Metal rings or bars are cemented into the mouse’s headcap, allowing them to be secured in the setup. Servos control the extension, retraction, and rotation of a spout head, each of which can dispense a different liquid. Liquids are gravity fed with flow controlled by a solenoid, and licks are detected using a capacitive sensor on the metal lick spout. All components are controlled by an Arduino.

Mice were water restricted to 90% of ad libitum body weight and were initially trained to lick by providing 20% sucrose in all 5 spouts. Mice experienced 2 days with short training sessions (30 trials, 3 s of access per trial) and 1 full length training session (60 trials).

Following training, mice experienced 5 sessions (one per day) with one dry spout and one spout each dispensing water, 10% sucrose, 20% sucrose, or 20 mM saccharin. The spout order was randomized across days. Each session consisted of 60 trials (12 per spout) presented in a pseudorandom order with a variable 15-20 s ITI. During each trial one spout extended and was available for a 3 second lick access period. Each lick triggered delivery of ∼1.5 µl solution.

After mice were tested under water restriction, they were returned to ad libitum water for at least 3 days, and then were food restricted to 85% of ad libitum body weight and experienced 5 additional multispout sessions.

For head-fixed TeTox experiments, control mice expressed AAV-FLEX-YFP, or were GCaMP-expressing mice from Figure 1, run in the same cohorts with TeTox mice.

### Body composition analysis and indirect calorimetry

Body composition analysis of body fat mass and lean mass was determined using quantitative magnetic resonance spectroscopy (EchoMRI 3-in-1; Echo MRI)^37^ at the University of Washington Nutrition Obesity Research Center (NORC) Energy Balance Core. For indirect calorimetry studies, also performed at the NORC Core, mice were acclimated to metabolic cages after which energy expenditure was measured using a computer-controlled indirect calorimetry system (Promethion Core, Sable Systems, NV) as described previously^38^. For each animal, O_2_ consumption and CO_2_ production were measured for 1-min at 5-min intervals. Respiratory quotient was calculated as the ratio of CO_2_ production to O_2_ consumption. Energy expenditure was calculated using the Weir equation^39^. Ambulatory activity was measured continuously, with consecutive adjacent infrared beam breaks in the x-, y-and z-axes scored as an activity count that was recorded every 5 or 10 min. Data acquisition and processing were coordinated by PromethionLive and Sable Systems Macro Interpreter (version 23.6) using an analysis script detailing all aspects of data transformation.

### Open field locomotion

Mice were placed into a large rectangular arena (80 x 50 x 40 cm L x W x H) and allowed to freely explore for 5 minutes while photometry signals were recorded.

### Core body temperature measurements

Body temperature was measured using a digital probe (Digi-Sense) inserted 2 cm into the rectum for 10 s.

### Nestlet shredding

Singly housed mice were placed into a clean cage with a fresh nestlet. Photos of the shredded nestlet were taken 24 hours later and were scored on a 5 point scale^24^ by two investigators blind to group. The assay was repeated twice per mouse and scores were averaged to generate a single composite score per mouse.

### High fat diet and Ensure

Standard chow used in this study was 4.5% kcal fat (LabDiet 5053). To measure weight gain on a solid high fat diet mice were given ad libitum access to a 60% kcal diet (Research Diets D12492). To measure weight gain on a liquid diet, chow was removed and mice were given ad libitum access to vanilla Ensure via a test tube sipper, with water available via a second sipper.

### Operant lever pressing

Fiber photometry mice were food restricted to 85% of ad libitum body weight and placed into an operant chamber (MedAssociates) and received 1 pre-training session in which 20 non-contingent pellets (20 mg Purified Dustless Precision Pellets, Bio-Serv) were dispensed with a variable 90 s ITI. Next, mice experienced 5 days of delayed cue operant training, in which both levers extended but only one was active. A press on the active lever led to a 3 s delay, followed by a delivery of a 3 s compound cue (lever light plus 3 kHz tone), followed by pellet delivery. The house light extinguished after each rewarded press and came back on after a 12.5 s ITI to signal the start of a new trial. Training sessions lasted 1 hour. Three photometry mice earned fewer than 2 pellets on day 1 of training, and for these mice the photometry data from day 2 was treated as day 1 for analysis. One of these mice earned fewer than 10 pellets on day 2 of training (treated as day 1), and was excluded from the comparison of first and last five trials for that day. For tests of untethered TeTox versus control mice, animals did not receive a pre-training session.

### Sucrose preference

Water was removed from the animal’s home cage for 6 hours prior to the start of the test, while ad libitum chow remained available. One sipper containing water and one sipper containing 1% sucrose were then added to the cage, and the amount of each liquid consumed was measured at 1, 2, and 16 hour time points.

### FED3 devices

A FED3 device^22^ loaded with Grain-Based Dustless Precision Pellets (20 mg, Bio-Serv) was placed in the home cage with a singly housed mouse. All chow was removed and water was available ad libitum. Mice were run on a free feeding program for 48 hours, in which a new pellet was dispensed into the hopper every time the previous pellet was removed. Mice were monitored every 24 hours and those that did not engage with the device were given a small amount of supplemental chow if needed to ensure they did not drop below 85% of ad libitum weight. Mice then experienced 24 hours each of an FR1, FR3, and FR5 paradigm, in which the required number of nose pokes in the active port were required to dispense a pellet. Nose pokes and pellet retrieval times were stored on an SD card for offline analysis.

### Fast/refeed in open field

Mice were fasted for 24 hours prior to being placed in a square open field arena (42 x 42 x 30 L x W x H) with a single chow pellet affixed via rubber band to a petri dish in the middle of the arena. Mice were video recorded during a 20 minute trial for analysis with Ethovision software (Noldus). The change in weight of the pellet (i.e. food consumed) was also recorded.

### Pavlovian fear conditioning

Day 1 consisted of a pre-conditioning test (pretest) in the morning and a first fear conditioning session (cond. 1) in the afternoon. Day 2 consisted of a post-conditioning retention test (probe 1) in the morning and a second fear conditioning session (cond. 2) in the afternoon. Day 3 consisted only of a post-conditioning retention test (probe 2) in the morning. All sessions were conducted in operant boxes (MedAssociates), and two contexts were created: one for pretest and probe sessions (Context A) and another for conditioning sessions (Context B). In context A, the boxes were fitted with solid white plastic walls that covered all surfaces, including the shock grids, and scented with a 1% acetic acid solution. In context B, the surface coverings were removed and the box was scented with 70% ethanol. Mice were placed in the operant boxes and underwent a 2 min habituation period, after which an auditory CS (10 kHz, 9 s duration, 60 s ITI) was presented 5 times for pretest and probe sessions and 10 times for conditioning sessions. During conditioning sessions, the CS terminated with a 0.5 s footshock (0.3 mA).

### Data Processing and Statistical Analysis

#### General statistics

Statistical tests were performed using Python or Prism 11 (GraphPad). The Geisser-Greenhouse correction was used to correct for unequal variability of differences in repeated-measures ANOVA tests.

#### Fiber photometry analysis

Processing of fiber photometry signals was performed using custom Python code adapted from^40^. Briefly, 465 and 405 signals were downsampled 100x and each was independently fit to a double exponential curve. The curve was subtracted to account for bleaching, and the GCaMP signal was motion corrected using a linear regression of the correlation between the 465 and 405 signals. For headfixed recordings the isosbestic channel was not used and traces were corrected for bleaching only. The corrected signal surrounding each timestamped event was extracted, and traces were z-scored to the indicated baseline on a per trial basis, unless otherwise noted. All z-scored trials for a given animal were averaged to generate a per-animal mean, unless otherwise noted.

### Linear mixed effects model

Data from all multispout sessions under water restriction and food restriction were corrected for bleaching and concatenated. A combined baseline for each mouse was generated by extracting the 5 s prior to each trial across all days, and z scores were calculated using this baseline. A single average z score for 0-10 s following spout extension was then calculated for each trial and used in the model.

The model was fit with statsmodels (Python) using the mixedlm function, with maximum likelihood (ML) estimation. Fixed effects were: restriction state, solution, previous solution, session number, trial number within the session, and number of licks per trial. Interaction terms (lick number x solution, restriction state x lick number, and restriction state x solution) were added where indicated. Solution was treated as a categorical variable and compared to the dry spout reference value, while the effect of previous solution for each trial was compared to the mean across all previous solution categories. For restriction state, water restricted was treated as the reference value. Session, trial number, and lick count were grand-mean centered. Random intercepts were fit per mouse, and a mouse×restriction×session variance component was included to allow session-specific baselines within each animal and restriction block. Predictor importance was assessed with drop-one ML comparisons (ΔAIC, likelihood ratio test). Fixed-effect significance was assessed with Wald z-tests (95% confidence intervals) from the model fit. For centered continuous predictors (session, trial, lick count), effects were expressed per one standard deviation of the predictor to facilitate comparison of magnitude.

### Velocity correlation

Photometry signals were corrected for bleaching and motion as described above, then were low-pass filtered (Butterworth, 1 Hz, zero phase) and z-scored to the session mean and standard deviation. Video recordings were tracked using Ethovision and center-point velocity was smoothed and mapped onto photometry timestamps. The Pearson’s r between the velocity and z-scored photometry was calculated per mouse, and values were Fisher z-transformed to calculate the mean and CI, then back-transformed for plotting. 1,000 random circular permutations of the velocity trace relative to the photometry trace were calculated per mouse and averaged to generate one null mean z per permutation for comparison with the observed mean.

### Atlas alignment and terminal mapping quantification

Free floating 50 µm sections were collected from brains expressing synaptophysin-GFP in LH-Nts neurons. Every third section was stained with anti-GFP antibody (AbCam 13970), mounted onto glass slides, and imaged at 10x with a VS200 slide scanner microscope (Evident Scientific). Each section image was aligned to the Allen Brain Atlas using the Aligning Big Brains & Atlases (ABBA)^41^ Fiji plugin in conjunction with QuPath^42^ image analysis software. The Subcellular Detection tool in QuPath was used to identify GFP puncta. The integrated fluorescence intensity of each puncta was calculated as the mean puncta intensity multiplied by the puncta area. The summed integrated intensity of all puncta within a region was then divided by the total area of the region to generate an integrated intensity per square micron measurements. To account for differences in viral injection or staining between animals, these values were normalized within animal to the combined integrated intensity per micron of all regions. Regions with normalized intensities greater than 5% of the LH intensity were plotted.

## Code Availability

All code used for data analysis is available at github.com/sodenlab/Sumarli-et-al

## Acknowledgements

We thank Selena Schattauer and Ella Kirwan for viral production in the Molecular Genetics Resource Core. This work was supported by NIH grants R01 DA054924 (M.E.S), K99 DA059612 and F32 DA054719 (A.G.), the University of Washington Center of Excellence in Opioid Addiction Research (P30 DA048736), a NIH High-End Instrumentation Grant S10 OD036208 (G.J.M.), the NIDDK-funded Nutrition Obesity Research Center (P30 DK035816) at the University of Washington, and a University of Washington Medicine Diabetes Institute Pilot and Feasibility Award to M.E.S.

## Author contributions

Conceptualization and interpretation: M.E.S., G.J.M, A.G, G.D.S. Data collection: D.S., M.C.L, G.O.D., K.W.S., S.M., C.L.B., E.V., M.K. Data analysis and visualization: D.S., K.W.S., C.L.B., M.E.S. Analysis code: M.E.S., D.S. Writing: original draft: M.E.S. Writing: review and editing: all authors.

## Declaration of interests

The authors declare no competing interests.

## Notes

### Competing Interest Statement

The authors have declared no competing interest.

