## Supplemental Material for "Multidimensional control of ingestive behavior by lateral hypothalamic neurotensin neurons"

### Supplemental Figures

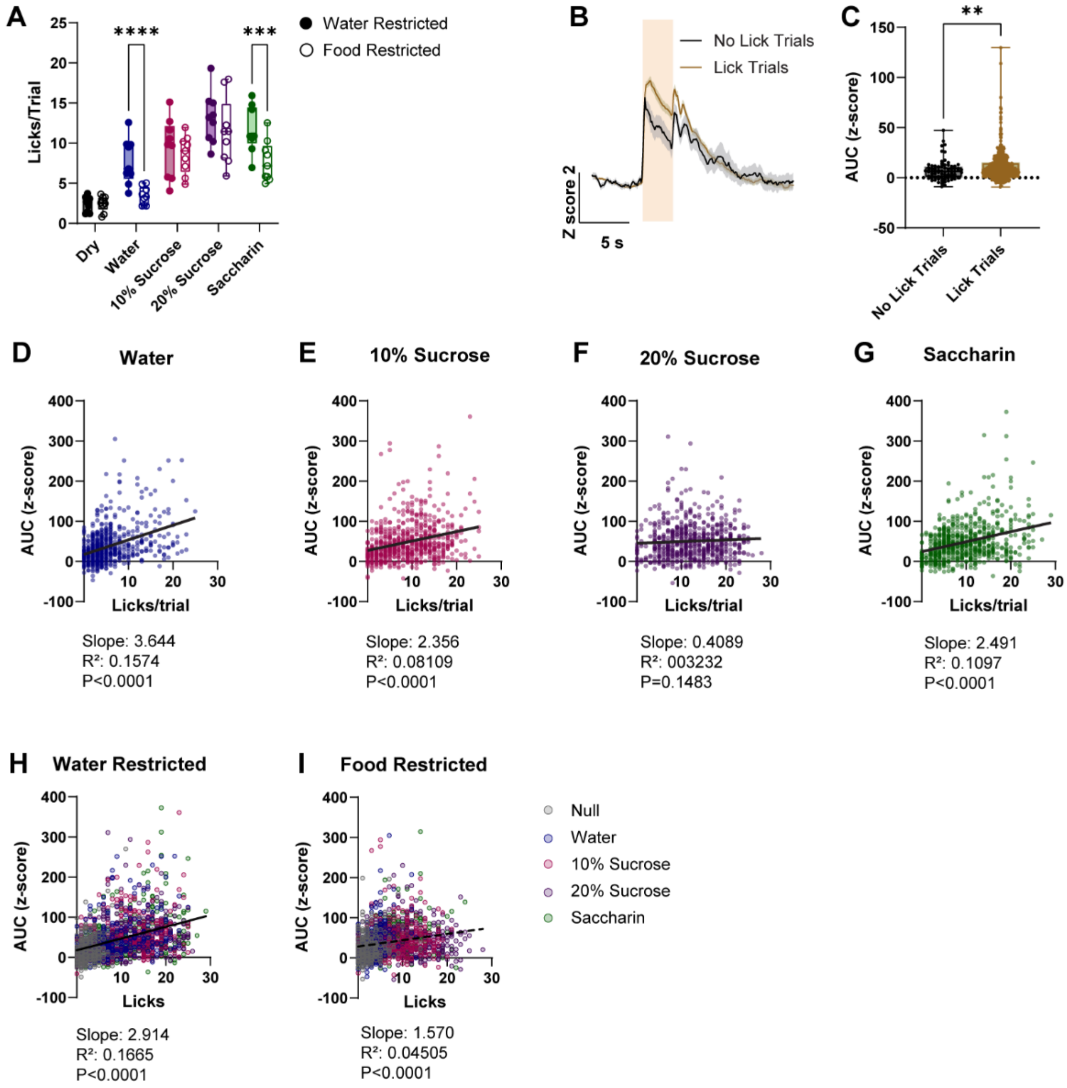


**Supplemental Figure 1.**

**A)** Licks per trial, comparing water and food restricted conditions (N=9 mice, $F_{(4, 32)}$=61.31 P<0.001. Sidak's multiple comparisons ***P<0.001, ****P<0.0001). **B)** Average GCaMP photometry traces for all dry spout trials (days 3-5 of training, both food and water restricted conditions), divided into trials with zero licks and trials with 1 or more licks. Shaded region = time of spout extension. **C)** AUC of the photometry z score for 0-10 s following spout extension in no lick and lick dry spout trials (76 no lick trials, 572 lick trials, Unpaired t test, P=0.0033). **D-G)** Linear regression of the number of licks and photometry signal AUC of individual trials for the indicated solution (days 3-5 of training, both food and water restricted conditions. **H-I)** Linear regression of the number of licks and photometry signal AUC of individual trials for the indicated solution under water restriction (H) or food restriction (I).

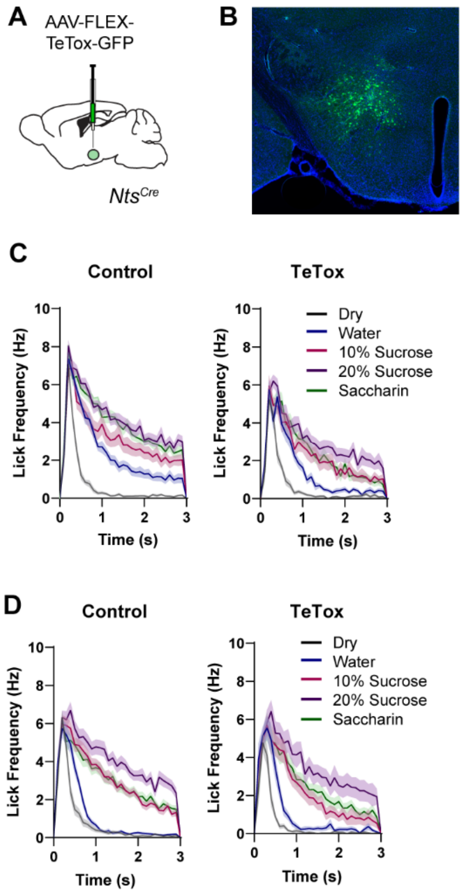


**Supplemental Figure 2.**

**A)** Schematic of AAV-FLEX-TeTox-GFP injection into the LH of $Nts^{Cre}$ mice. **B)** Example histology image of TeTox-GFP in the LH. **C)** Lick frequency for each solution across the 3 s access period for mice under water restriction. **D)** Lick frequency for each solution across the 3 s access period for mice under food restriction.

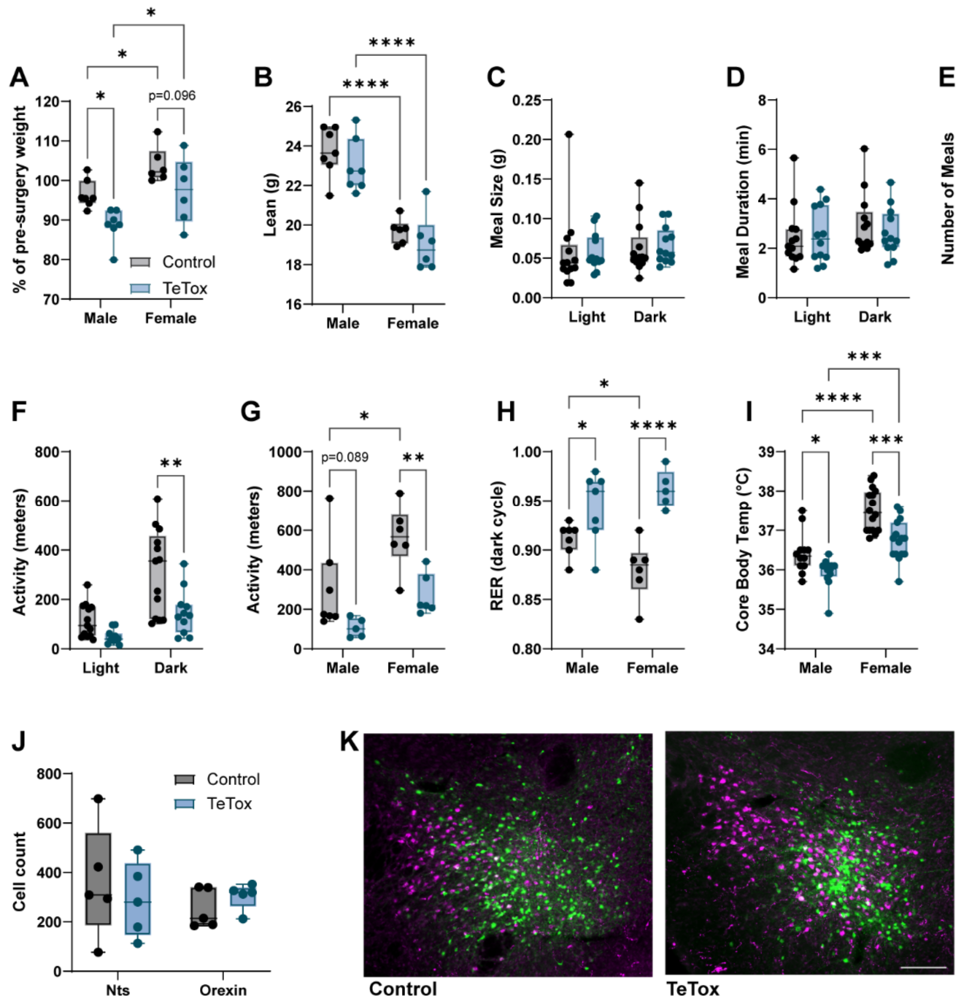


**Supplemental Figure 3.**

**A)** Body weight as a percentage of starting weight before viral injection (N=7 male, 6 female mice per group; 2-way ANOVA, effect of sex: $F_{(1, 22)}$=14.47, P=0.0010, effect of virus: $F_{(1, 22)}$=11.44, P=0.0027, Sidak's multiple comparisons *P<0.05). **B)** Lean mass (2-way ANOVA, effect of sex: $F_{(1, 22)}$ = 67.24, P<0.0001, Sidak's multiple comparisons ****P<0.0001). **C)** Grams consumed per meal during the light and dark cycle. **D)** Meal duration during the light and dark cycle. **E)** Number of meals during the light and dark cycle. **F)** Total activity during the light and dark cycle (2-way RM ANOVA, $F_{(1, 22)}$ = 5.892, P=0.0239, Sidak's multiple comparisons **P<0.01). **G)** 24 hour activity separated by sex (2-way ANOVA, effect of sex: $F_{(1, 20)}$=10.43, P=0.0042, effect of virus: $F_{(1, 20)}$=14.09, P=0.0012, Sidak's multiple comparisons *P<0.05, **P<0.01). **H)** RER during the dark cycle separated by sex (2-way ANOVA, $F_{(1, 21)}$ = 5.069, P=0.0352, Sidak's multiple comparisons *P<0.05, ****P<0.0001). **I)** Core body temperature separated by sex (N=12 male control, 16 female control, 12 male TeTox, 15 female TeTox, 2-way ANOVA effect of sex: $F_{(1, 51)}$=49.53, P<0.0001, effect of virus: $F_{(1, 51)}$= 21.58, P<0.0001, Sidak's multiple comparisons *P<0.05, ***P<0.001, ****P<0.0001). **J)** Number of virally labeled Nts cells and immunostained orexin cells (N=5 mice/group, 3 sections counted per mouse). **K)**

Example images of control and TeTox-expressing cells (green) and orexin staining (magenta). Scale bar = 200 µm.

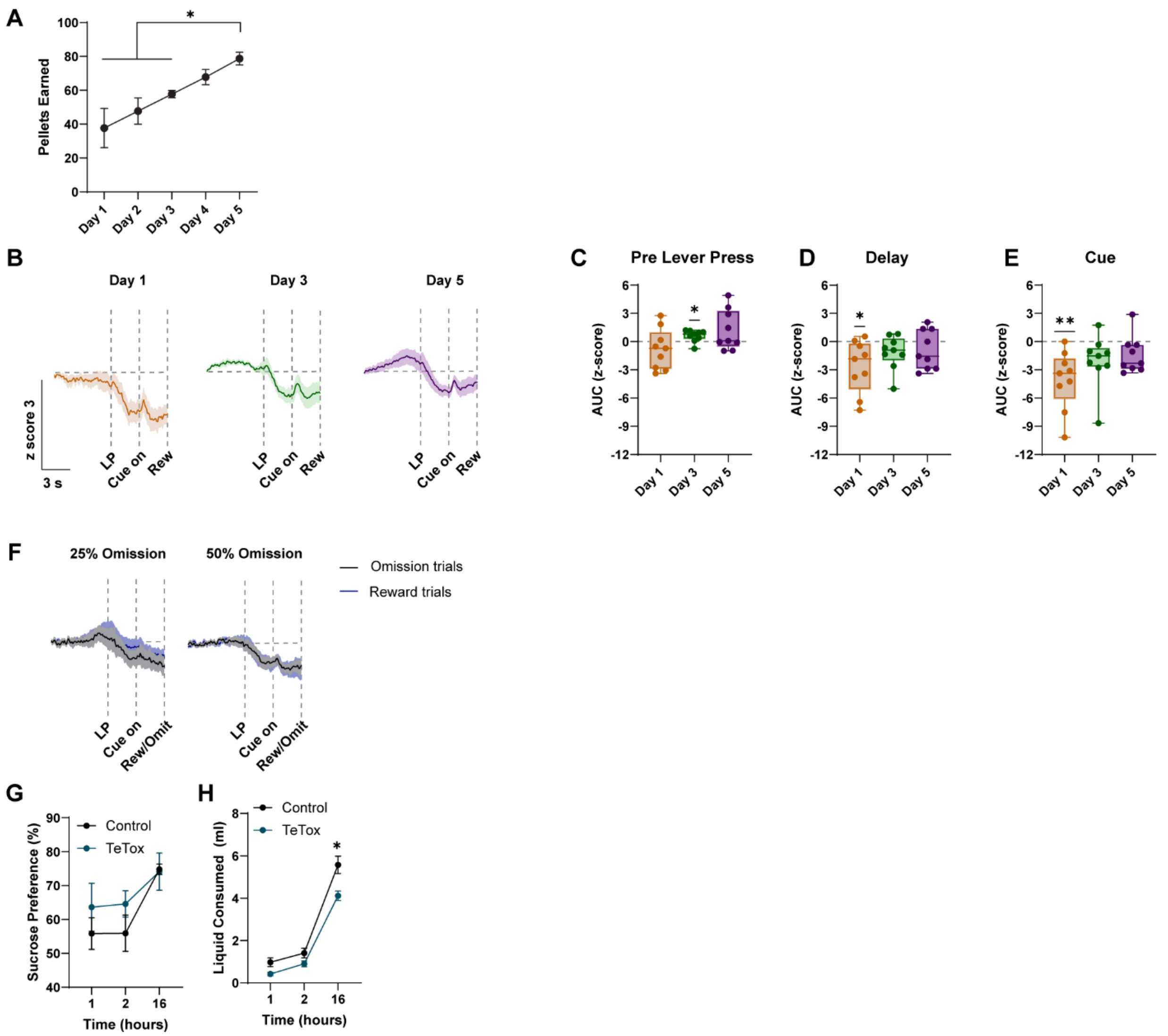


**Supplemental Figure 4.**

**A)** Number of pellets earned across days in operant task (N=9 mice, One-way RM ANOVA $F_{(1.486,\ 11.89)}$=6.142, P=0.0203, Tukey's multiple comparisons *P<0.05). **B)** Average z-scored photometry trace aligned to rewarded lever press on indicated days. A 4 s baseline period beginning 10 s prior to the lever press was used to calculate the z score (N=9 mice). **C-E)** AUC of the z score during the 3 s prior to the lever press, during the 3 s delay, or during the 3 s cue period. (One-sample t test comparing to zero, *P<0.05, **P<0.01). **F)** Average z-scored photometry trace aligned to the lever press for rewarded and omitted trials on indicated days. **G)** Sucrose preference at indicated time points (amount of 1% sucrose consumed as a percentage of total liquid consumed; N=12 control 9 TeTox, 2-way RM ANOVA, Effect of time: $F_{(1.553,\ 29.50)}$=8.399, P=0.0026). **H)** Liquid consumed during the sucrose preference test at the indicated time points (2-way RM ANOVA $F_{(1.197,\ 22.74)}$=6.670, P=0.0130, Sidak's multiple comparisons *P<0.05).

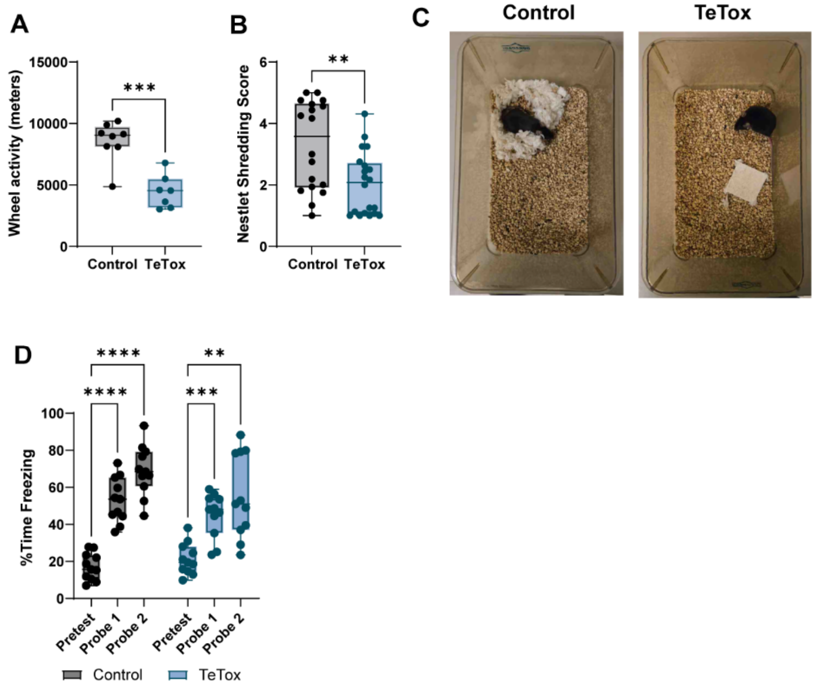


**Supplemental Figure 5.**

**A)** 24 hour running wheel activity (unpaired t test P=0.0002). **B)** Nestlet shredding score after 24 hours, average of 2 investigators scoring each of 2 tests per mouse (N=18 control, 20 TeTox, unpaired t test P=0.0046). **C)** Example nestlet images. **D)** Percent time freezing during cue presentation during the pretest and probe tests (N=11/group, 2-way RM ANOVA, $F_{(1.352, 27.03)}$=4.029, P=0.0437, Sidak's multiple comparisons **P<0.01, ***P<0.001, ****P<0.0001).
